# Transepithelial Electrical Impedance Increase Following Porous Substrate Electroporation Enables Label-Free Delivery

**DOI:** 10.1101/2023.10.17.562630

**Authors:** Justin R. Brooks, Tyler C. Heiman, Sawyer R. Lorenzen, Ikhlaas Mungloo, Siamak Mirfendereski, Jae Sung Park, Ruiguo Yang

**Affiliations:** Department of Mechanical and Materials Engineering, University of Nebraska-Lincoln, Lincoln, NE, 68588, USA; Nebraska Center for Integrated Biomolecular Communications, University of Nebraska-Lincoln, Lincoln, NE, 68588, USA

**Author notes:** Corresponding author: R.Y.

**Keywords:** porous substrate electroporation, transepithelial electrical impedance, label-free delivery

## Abstract

Porous substrate electroporation (PSEP) is a promising new method for intracellular delivery, yet fundamentals of the PSEP delivery process are not well understood, partly because most PSEP studies rely solely on imaging for evaluating delivery. Although effective, imaging alone limits understanding of intermediate processes leading to delivery. PSEP is an electrical process, so electrical impedance measurements naturally complement imaging for PSEP characterization. In this study, we developed a device capable of measuring impedance and performing PSEP and we monitored changes in transepithelial electrical impedance (TEEI). Our measurements show TEEI increases following PSEP, unlike other electroporation methods. We then demonstrated how cell culture conditions and electrical waveforms influence this response. More importantly, we correlated TEEI response features with viability and delivery efficiency, allowing prediction of outcomes without fluorescent cargo, imaging, or image processing. This label-free delivery also allows improved temporal resolution of transient processes following PSEP, which we expect will aid PSEP optimization for new cell types and cargos.

**TEASER:** Electrical impedance measurements were used to understand delivery and cellular response after porous substrate electroporation.

## INTRODUCTION

Electroporation is one of the most common methods of intracellular delivery and involves temporary permeabilization of the cell membrane using an electric field. A subset of electroporation known as porous substrate electroporation (PSEP) has been gaining traction as a promising new delivery method (*1–5*). PSEP involves culturing adherent cells on a porous substrate that focuses the electric field to discrete portions of the basal cell membrane (*1, 6*). PSEP has several unique attributes that make it a promising delivery method, including reduced stress from delivering to cells in an adherent state, the ability to perform long-term *in situ* delivery due to adherence, reduced electric field exposure from focusing the electric field through the porous substrate channels (*1*), and more consistent electric field exposure due to more uniform positioning of cells relative to the electrodes (*5*). Recent studies have utilized these advantages for applications such as efficient genetic editing (*2*) and the extraction of intracellular molecules (*3*). Despite these applications, PSEP research is limited by its reliance on fluorescent microscopy as the only method for assessing delivery outcomes. While fluorescent tags are an effective indicator at the end of the delivery process, intermediate stages such as cargo transport through the substrate channels, cell membrane permeabilization, cargo delivery into the cytoplasm, and membrane resealing are difficult to assess using fluorescent microscopy alone (*7*). In addition, there are many variables that influence the success of PSEP delivery, including characteristics of the delivered molecule, differences between cell types, waveform parameters, electrolytes, substrates, electrodes, and so forth (*4*). Since PSEP assessment is dependent on fluorescent imaging, or subsequent blotting or sequencing, testing each combination of experimental parameters requires time-consuming image collection and processing, as well as expensive reagents (*8*).

These challenges can be mitigated by augmenting PSEP assessment with impedance measurements during and after electroporation. The electrical impedance of the cell membrane and porous substrate channels contains an abundance of information about the cell state and the transport of molecular cargos during PSEP, with untapped potential for understanding the PSEP process. We previously demonstrated how impedance measurements can be used prior to PSEP to better understand the influence of different system parameters and estimate the electrical waveforms needed for delivery (*9*). Impedance can also be measured intermittently throughout delivery to provide additional information on the intermediate stages of the process, and it may even be possible to optimize PSEP using only impedance measurements, a method known as label-free delivery (*7*). These impedance measurements during and after electroporation have already been utilized with many different electroporation methods (*10, 11*), but mostly for assessing membrane permeabilization and to our knowledge, have not been reported for PSEP. More specifically, a subset of impedance measurements known as transepithelial electrical impedance (TEEI) should be utilized with PSEP. TEEI is the impedance of the cell monolayer multiplied by the area of the culture surface. TEEI is similar to the more commonly used transepithelial electrical resistance (TEER), with the distinction that TEER is measured at a single low frequency, whereas TEEI is measured at multiple higher frequencies (*12*). TEEI is well suited for implementation with PSEP because it can leverage existing TEEI research describing how cell behavior influences the electrical system (*13, 14*). Moreover, both PSEP and TEEI utilize electrodes on both sides of a cell monolayer cultured on a porous substrate. Despite these potential advantages, TEEI has not yet been demonstrated before and after PSEP due to the lack of a device capable of measuring TEEI while performing PSEP.

In this study, we investigated whether we could estimate the result of PSEP by monitoring TEEI following electroporation. Using a new platform that combines cell culture inserts and customized electronics, we observed increases in TEEI following PSEP. We conducted comprehensive experiments to characterize TEEI responses across different cell types, under varied cell culture conditions, and with different electrical waveform parameters. More importantly, we demonstrated how features of the TEEI response are correlated with delivery efficiency and cell viability readings, indicating TEEI monitoring can assist label-free delivery for PSEP. In addition to measuring TEEI following PSEP, this study shows how TEEI measurements following electroporation can be used to better understand the intermediate processes of PSEP.

## RESULTS

### TEEI measurement before and after electroporation

We developed an integrated device for PSEP and TEEI measurement using customized electronics and commercialized cell culture inserts (Figure 1A-C). Our device utilizes electrode arrays capable of measuring and electroporating 6 samples in parallel (Figure 1C). Each array consists of cell culture wells, 3D printed using biocompatible resin, and upper and lower printed circuit boards (PCBs) for the electrodes. The upper and lower PCBs each contain distinct voltage measurement and current injection electrodes (Figure 1B), allowing for 4-electrode impedance measurements which improve measurement accuracy by removing electrode-electrolyte interface impedances from the measurement (*15*). We verified our device could measure TEER values with comparable accuracy to widely used systems (Figure S1C) and record impedances across a range of frequencies relevant to PSEP with the same accuracy as an LCR meter (Figure S1D-F, Table S1). Our device also utilizes chamber electrodes that provide improved TEEI measurement consistency compared to chopstick electrodes by ensuring consistent and symmetrical electrode placement (*15*). Furthermore, our device incorporates commercially available cell culture inserts, which were recently used for PSEP for the first time (*16*), thus standardizing the PSEP process.

**Figure 1.**
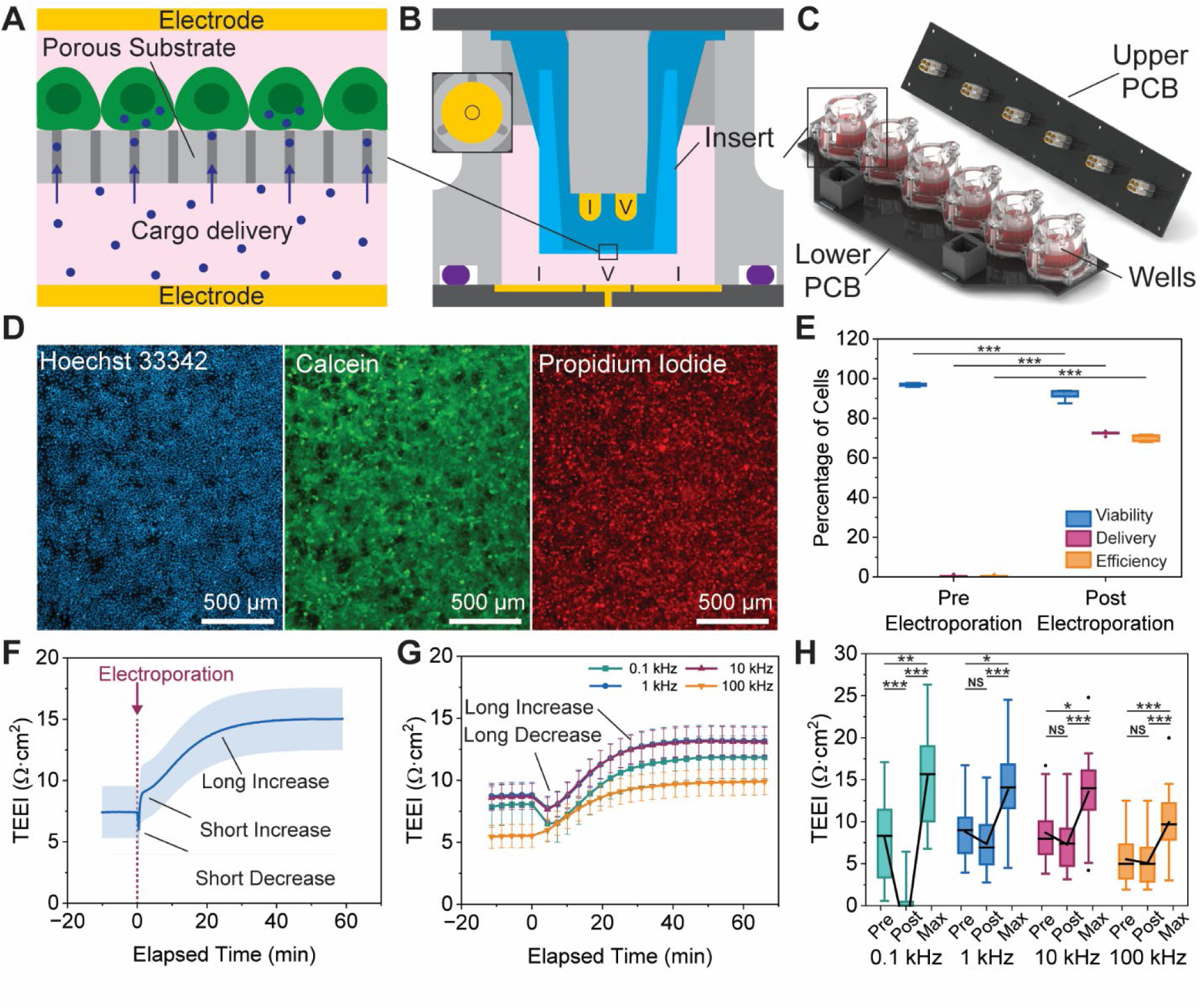
PSEP induces a TEEI increase. **A.** Simplified depiction of PSEP, with cargo molecules passing through substrate channels and into adherent cells. **B.** Cross-sectional view of a well in the working configuration. The insert is shown in blue, cell culture media in pink, electrodes in gold, well walls and pin cover in gray, PCBs in black, and O-ring in purple. Current injection electrodes are marked “I” and voltage measurement electrodes are marked “V”. The inset shows a top-down view of the concentric lower electrodes. All dimensions are to scale except the substrate and lower electrodes which were thickened for visibility. **C.** Rendering of an electrode array with the upper PCB removed. **D.** Fluorescent microscope images immediately following electroporation stained with Hoechst 33342, calcein, and PI to show electroporation occurred. **E.** Quantification of viability, PI delivery, and PI delivery efficiency from fluorescent images before and after electroporation. **F.** Post-EP TEEI response measured at 1 kHz at our device’s maximum sample rate. **G.** Post-EP TEEI response measured simultaneously at multiple frequencies at a reduced sample rate. **H.** TEEI comparison at multiple frequencies and multiple time points. For the box plots in E (n = 6) and H (n = 21), the shaded area represents the interquartile range (IQR), the error bars represent 1.5 times the IQR, the horizontal line is the median, the lines connecting between time points are the averages, and the diamonds are outliers that are located beyond the error bars. For the time plots in F (n = 15) and G (n = 21), error bars represent the SEM. For the box plot in H, “Pre” is defined as the TEEI immediately before PSEP, “Post” is defined as the TEEI immediately after PSEP or at the lowest point if applicable, and “Max” is defined as the maximum TEEI after PSEP.

Before measuring changes in impedance following electroporation, we needed to find waveform parameters that would cause electroporation in our system. We found waveforms consisting of 30 V, 1 ms pulse duration, 20 Hz pulse frequency, and 200 pulses yielded an average of 92% viability, 72% propidium iodide (PI) delivery, and 70% delivery efficiency (the percentage of cells both living and delivered to) (Figure 1D-E). These waveform parameters are similar to parameters used in other PSEP studies (*3, 4, 6, 17*). We then applied these waveforms and measured the impedance before and after electroporation. It is worth noting that samples were incubated for 1h prior to impedance measurement to allow for thermal equilibration, followed by 10 minutes of impedance measurement at 1 kHz to establish baseline impedance, electroporated for 10 seconds, and finally 1h of additional impedance measurements (Figure 1F). 1 kHz was chosen for the measurements because at this frequency the cell monolayer has one of its largest contributions to the total system impedance (*14*).

We consistently observed a substantial increase in impedance following electroporation (Figure 1F). This response was only observed following electroporation in samples containing cells and consisted of an exponentially plateauing TEEI increase that achieved 95% of the increase after 31 minutes. The response is especially significant when considering its size relative to the initial TEEI. The initial TEEI was approximately 6.7 Ω·cm^2^ and the final TEEI following electroporation was approximately 14.0 Ω·cm^2^, more than doubling the baseline. We have observed a total of 4 distinct features to the post-electroporation (post-EP) response, including 1) a short decrease, 2) a short increase, 3) a long decrease, and 4) a long increase, although not all 4 features are observed in all samples. Of these 4 features, the short decrease and short increase are observed in both samples with and without cells (Figure S2A) and are thought to be artifacts due to electrode corrosion and polarization at the electrode-electrolyte interface following electroporation. When subtracting the samples without cells from the samples with cells during the TEEI calculation, these features are mostly reduced but are not always eliminated, likely due to slight variations in measurement.

The post-EP response was then measured at four frequencies (100 Hz, 1 kHz, 10 kHz, and 100 kHz) simultaneously to better understand the nature of the long decrease and long increase in impedance (Figure 1G-H). These frequencies were chosen because below 100 Hz the total impedance is heavily influenced by the electrode-electrolyte interface impedances, and above 100 kHz the cell monolayer impedance becomes very small (*14*). In addition, we measured all 6 samples in the electrode array simultaneously to allow for a higher throughput. This combination of multiple frequencies and parallel sample measurement meant that each sample would be measured once at the first frequency, then each sample would be measured once at the second frequency, and so on (as shown in Table S2). These changes reduced our temporal resolution, which can be seen when comparing Figure 1F to Figure 1G, but this temporal resolution was sufficient to capture the response and its principal features, the long decrease and long increase, due to their relatively slow change and long duration. It is also evident that Figure 1F does not have the long decrease but Figure 1G does. The long decrease is present in some samples measured in Figure 1F (Figure S2B), but not the majority and the existence of the long decrease may depend on slight variations from sample to sample. There are also differences in the post-EP response between frequencies. The largest percent increase occurs at 100 kHz and the largest percent decrease occurs at 100 Hz, which dissipates at higher frequencies.

### Influence of cell culture conditions on the post-EP TEEI response

After observing an increase in TEEI following PSEP, we first ruled out noncellular explanations for the increase such as temperature change, corrosion, or bubble formation (Figure S3). We then examined whether the response was influenced by factors that influence PSEP, including cell type, confluency, and fibronectin coating (Figure 2). For each experiment in Figure 2, the standard electroporation waveform parameters, impedance measurement frequencies, and post-EP fluorescent imaging (Figure 2, column iii) noted in the materials and methods section were used. A detailed description of the cell culture parameters used in each experiment shown in Figure 2 can be found in Table S3.

**Figure 2.**
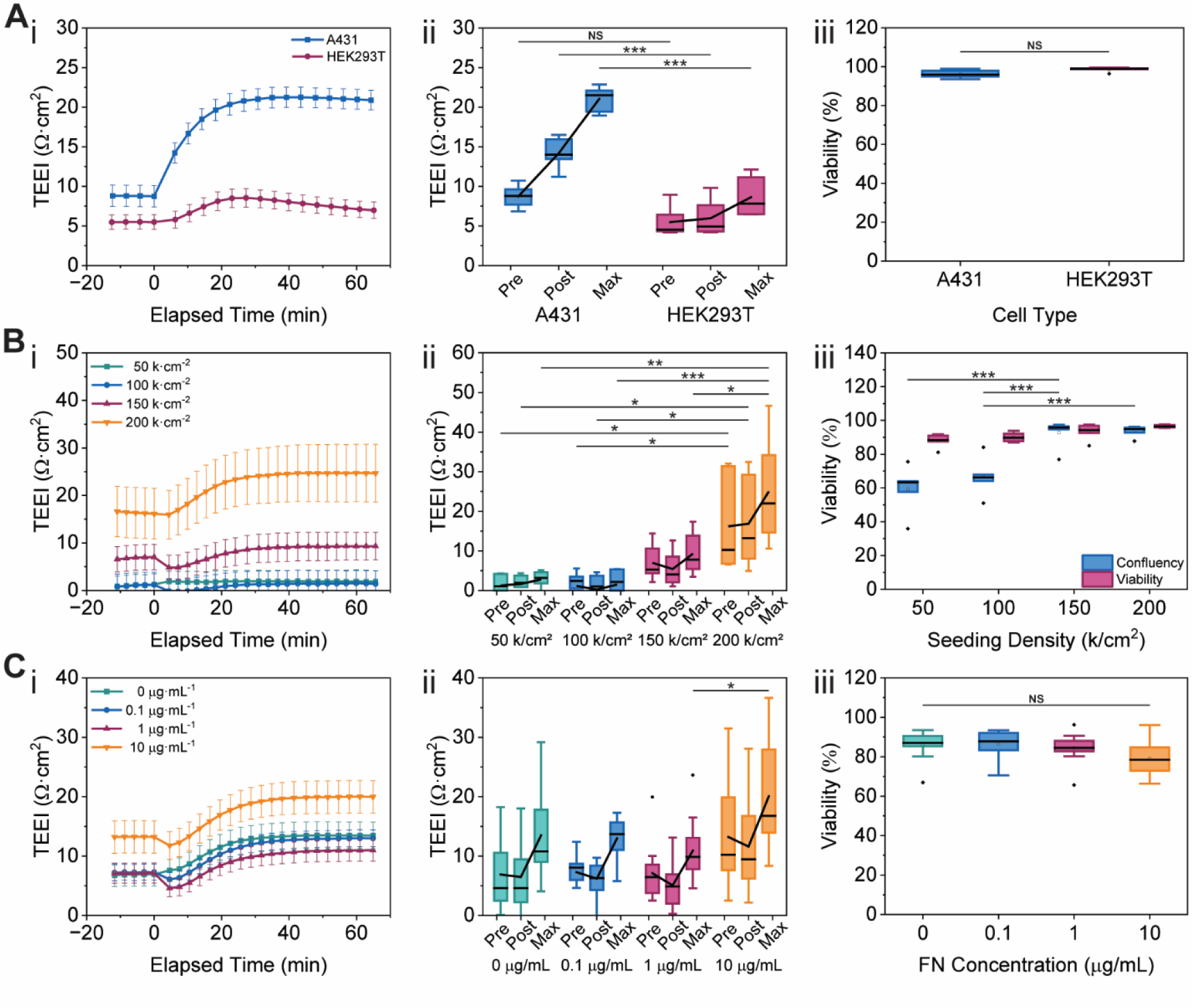
Cell culture conditions influence the post-EP TEEI response. A time plot (i), comparison of TEEI at the three main time points (ii), and comparison of viability at each condition (iii) are shown for each experiment. **A.** Post-EP TEEI response and corresponding viability at different seeding densities. **B.** Post-EP TEEI response and corresponding viability with different cell types. **C.** Post-EP TEEI response and corresponding viability at different fibronectin concentrations. For the time plots in i, error bars represent the SEM (n = 12). For the box plots in ii and iii, the shaded area represents the interquartile range (IQR), the error bars represent 1.5 times the IQR, the horizontal line is the median, the lines connecting between time points are the averages, and the diamonds are outliers that are located beyond the error bars (n = 12). For the box plots in ii, “Pre” is defined as the TEEI immediately before PSEP, “Post” is defined as the TEEI immediately after PSEP or at the lowest point if applicable, and “Max” is defined as the maximum TEEI after PSEP.

To understand whether the post-EP response is shared in other cell types, we measured the TEEI of A431 human epidermoid carcinoma cells and human embryonic kidney cells (HEK293T) (Figure 2A). We determined the seeding density that consistently produced approximately 100% confluency 12 hours after seeding for both cell types, with A431 seeded at 200,000 cells·cm^-2^, and HEK293T at 400,000 cells·cm^-2^. Although both cell types exhibited a post-EP TEEI increase, the responses were remarkably different. Prior to electroporation, the A431 cells started with a baseline of 8.8 Ω·cm^2^ and the HEK293T cells started with a baseline of 5.5 Ω·cm^2^. Following electroporation, the A431 TEEI increased much faster and to a much higher level than the HEK293T TEEI, and remained higher, whereas the HEK293T TEEI increased more slowly, quickly peaked, and began returning to baseline. The A431 TEEI increased to a peak of 21.3 Ω·cm^2^ after 43 minutes and remained at 20.9 Ω·cm^2^ after 65 minutes, whereas the HEK293T cells increased to a peak of 8.5 Ω·cm^2^ after 27 minutes and fell to 7.0 Ω·cm^2^ after 65 minutes. Despite the difference in post-EP response, there was no significant difference between the viability of the two cell types following electroporation. Differences in post-EP TEEI increase may be due to cell types responding to electroporation differently (*11*). This can be from physical differences, such as cell size and the strength of cell-cell junctions and cell-extracellular matrix (ECM) adhesions, or due to biological differences such as differences in susceptibility to transfection. Furthermore, different cell types have different TEEI measurements (*13*). The difference in response between the two cell types further suggests that the change in impedance is due to a biological response. From these experiments, A431 was chosen to be the main cell line for this study because it provided the largest change in TEEI, and therefore the best opportunity to understand the post-EP TEEI response. The larger TEEI response of the A431 cells may be due to the higher baseline TEEI of the A431 cells, perhaps due to tighter cell-cell junctions and cell-ECM adhesions prior to electroporation. The higher baseline TEEI may also increase the response by causing a larger percentage of the voltage applied to the system to drop across the cell monolayer.

We previously reported that PSEP delivery decreases at lower confluencies, which is thought to be due to a decrease in the cell monolayer impedance and corresponding decrease in voltage drop across the cell monolayer (*9*). To evaluate the influence of seeding density on the post pulse response, A431 cells were seeded at 50, 100, 150, and 200 thousand cells·cm^-2^ for 12 hours prior to electroporation (Figure 2B). 200,000 cells·cm^-2^ was sufficient to consistently achieve over 90% confluency 12 hours after seeding. Prior to electroporation, there was a difference in TEEI values between the seeding densities, with higher seeding densities having higher TEEI values, likely due to more confluent cell monolayers (*9*). Following electroporation, lower seeding densities experienced larger initial decreases in TEEI and smaller final increases in TEEI. Lower seeding densities also had lower viability, with cells on the edge of colonies more likely to be killed by the electroporation, perhaps due to increased electrical exposure. 200,000 cells·cm^-2^ was used as the standard seeding density for the remainder of this study to ensure confluent cell monolayers.

Increasing substrate coatings that promote cell adhesion, such as fibronectin and poly-L-lysine, has also been shown to increase PSEP delivery efficiency (*6*). The increased delivery efficiency is thought to be from stronger cell-substrate adhesion, which increases the voltage applied to the cell monolayer (*6*). We assessed the role of ECM coating concentration using concentrations of 0, 0.1, 1, and 10 μg/mL of fibronectin in phosphate-buffered saline (PBS) which were coated on the inserts for 3h in an incubator (Figure 2C). A431 cells were seeded and electroporated after 12h of culture. The 0, 0.1 and 1 μg/mL concentrations had similar TEEI values before and after electroporation, with larger initial TEEI decreases observed at the higher concentrations, perhaps due to stronger substrate attachment prior to electroporation, which yielded higher electric field exposure, resulting in damaged cell-cell junctions and cell-ECM adhesions and decreased TEEI. The 10 μg/mL concentration had a higher pre- and post-EP TEEI but a similar response to the lower fibronectin concentrations. The higher impedance for the 10 μg/mL concentration may be due to stronger cell-ECM adhesion and tighter cell-cell junctions. Although the A431 cells showed similar post-EP TEEI responses with and without fibronectin, fibronectin coatings were found to be critical to the existence of a post-EP response for HEK293T cells (Figure S4), perhaps due to inherent differences in cell-substrate attachment between the two cell types.

### Influence of electroporation waveform parameters on the post-EP TEEI response

After showing the post-EP TEEI increase is influenced by cell culture conditions, we investigated the influence of electrical waveform parameters on the post-EP response (Figure 3), since waveforms influence electroporation results, including viability, delivery, and delivery efficiency (*6, 11*). In this study we limited our analysis to unilevel square waveforms, which are the most widely used PSEP waveform, although waveforms such as exponential (*18*) and bilevel square waves (*19*) have also been used for PSEP. Unilevel square waveforms can be fully described with 6 parameters: voltage, pulse duration, pulse frequency, pulse number/pulse number per train, train frequency, and train number. Often only 1 train of pulses is used for PSEP, in which case only the first 4 parameters are necessary. We investigated the role of each of the first 4 parameters on the post-electroporation response (Figure 3A). For each waveform parameter investigated, all other waveform parameters were kept at standard values. For each experiment in Figure 3, the standard cell culture conditions, electroporation waveform parameters (30 V, 1 ms pulse duration, 20 Hz, 200 pulses), impedance measurement frequencies, and post-EP fluorescent imaging (Figure 3, column iii) noted in the materials and methods section were used. TEEI values in Figure 3 are reported as percentage change in TEEI from pre-pulse values since all samples had the same initial conditions. A detailed description of the waveform parameters used in each experiment shown in Figure 3 can be found in Table S4.

**Figure 3.**
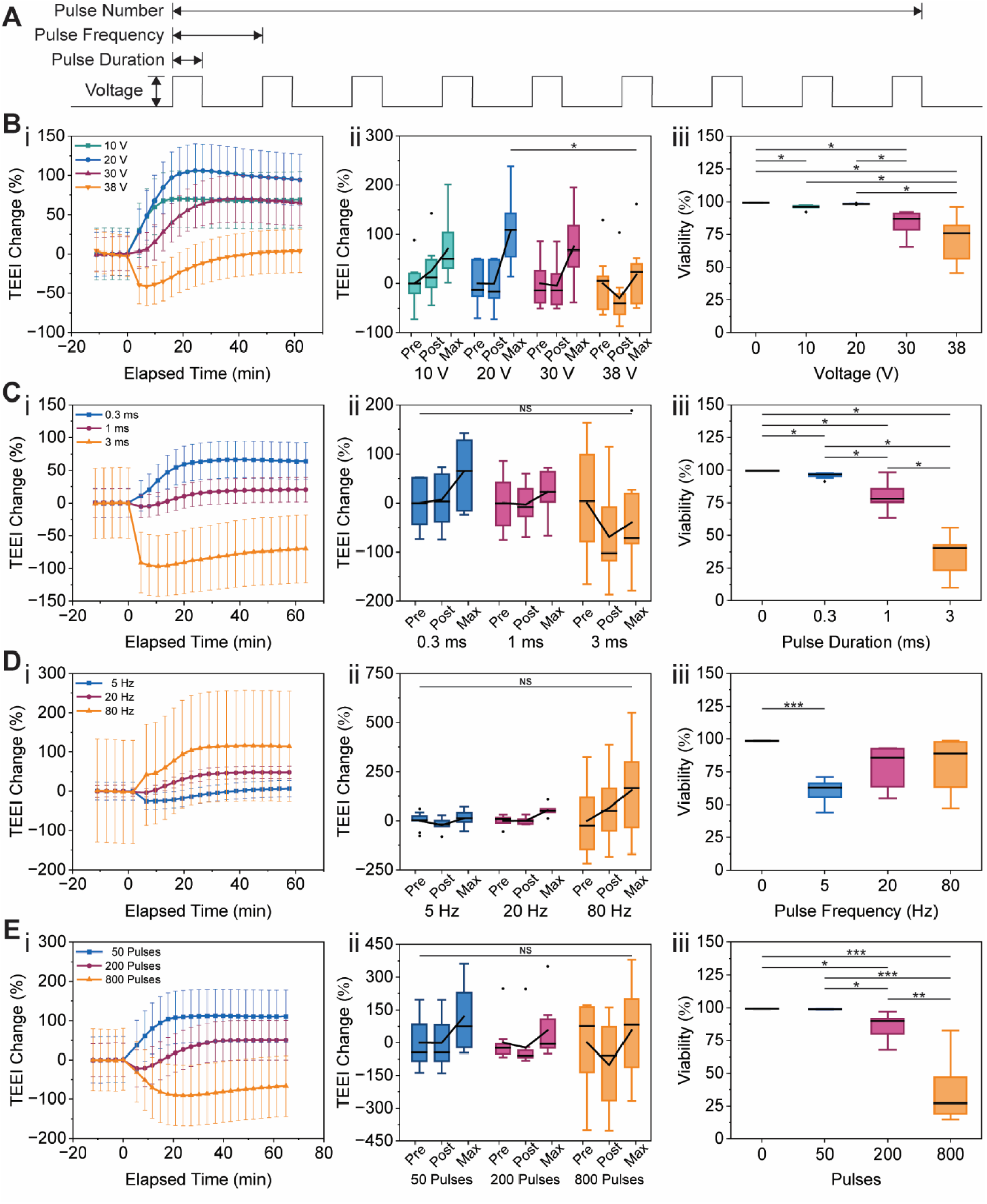
Electroporation waveform parameters influence the post-EP TEEI response. A time plot (i), comparison of TEEI at the three main time points (ii), and comparison of viability at each condition (iii) are shown for each experiment. **A.** Diagram showing the 6 parameters that describe an electroporation waveform. **B.** Post-EP TEEI response and corresponding viability at different voltages. **C.** Post-EP TEEI response and corresponding viability at different pulse durations. **D.** Post-EP TEEI response and corresponding viability at different pulse frequencies. **E.** Post-EP TEEI response and corresponding viability at different pulse numbers. For the time plots in i, error bars represent the SEM (n = 9). For the box plots in ii and iii, the shaded area represents the interquartile range (IQR), the error bars represent 1.5 times the IQR, the horizontal line is the median, the lines connecting between time points are the averages, and the diamonds are outliers that are located beyond the error bars. For the box plots in ii, “Pre” is defined as the TEEI immediately before PSEP, “Post” is defined as the TEEI immediately after PSEP or at the lowest point if applicable, and “Max” is defined as the maximum TEEI after PSEP.

The effect of voltage on the post-EP response was tested using voltages of 10, 20, 30, and 38 V (Figure 3B). 10 V to 20 V showed an increase in the post-EP response, but voltages above 20 V showed decreasing responses, with 38 V yielding a large decrease in TEEI before returning near baseline. Meanwhile, the viability assessment showed no effect on viability at 20 V and below, and an inverse relationship between voltage and viability above 20 V. Following assessment of the effect of voltage on the post-EP response, we tested the effect of pulse duration using 0.3, 1, and 3 ms pulses (Figure 3C). We observed an inverse correlation between pulse duration and post-EP response, with 3 ms showing a large decrease in TEEI. This trend corresponded to the viability assessment, which showed an inverse correlation between pulse duration and viability. The effect of pulse frequency on the post-EP response was tested using pulses at commonly used frequencies: 5, 20 and 80 Hz (*4*) (Figure 3D). 80 Hz showed the largest increase in TEEI without any initial decrease, followed by 20 Hz, and finally 5 Hz which showed an initial decrease followed by the smallest increase of the 3 pulse frequencies. This corresponded to quantification of cell viability, which showed a positive correlation between pulse frequency and cell viability. We expected the higher frequencies to have smaller post-EP increases and lower viability because the electroporation pulses are applied over a much shorter period, but it is possible that the prolonged application of pulses at the lower frequencies prevents the cells from recovering as quickly, which results in decreased viability. We also tested the effect of pulse number on the post-EP response using 50, 200, and 800 pulses (Figure 3E). Lower pulse numbers yielded higher post-EP increases, with 800 pulses showing a decrease in TEEI after electroporation. Lower pulse numbers also showed higher cell viability following electroporation, corresponding to the TEEI values.

Collectively, each of the waveform parameter experiments showed that increases in waveform parameters beyond our standard conditions resulted in decreases in TEEI response and cell viability, whereas decreases in waveform parameters resulted in increases in TEEI response and cell viability, suggesting the waveform parameters we chose as our standard are above the optimal parameters for inducing a TEEI increase and maintaining cell viability. This suggests that TEEI measurements may be used as an indicator of viability to optimize PSEP waveform selection.

### Post-EP TEEI enables label-free delivery

To understand the relationship between changes in the TEEI response and delivery metrics such as viability, delivery, and delivery efficiency, we first considered the general shape of the TEEI response at different voltages (Figure 4A-C). As mentioned, there are two primary features in the TEEI responses: a period of decreasing TEEI and a period of increasing TEEI. We quantified the increase and decrease in TEEI for all voltages, with the increase quantified as the increase from the minimum TEEI rather than the increase from baseline because in some instances (Figure 4C) measuring from baseline neglects most of the increase that has occurred. When the TEEI increase and decrease are plotted for each voltage, the TEEI increase grows parabolically to an optimal voltage before reducing, whereas the TEEI decrease grows exponentially with an increase in voltage (Figure 4E). Although the comparisons in Figure 4 are primarily focused on voltage, the same general trends are seen with other increases in electrical energy, such as increasing pulse duration or pulse number. These trends are similar to trends in delivery efficiency and cell death when increasing electrical exposure, where delivery efficiency is maximized at an optimal condition and cell death increases with electrical exposure (Figure 4F). Due to the similarities between the TEEI increase from minimum and delivery efficiency, and the TEEI decrease and cell death, we sought to determine whether the impedance response features were correlated to delivery outcomes.

**Figure 4.**
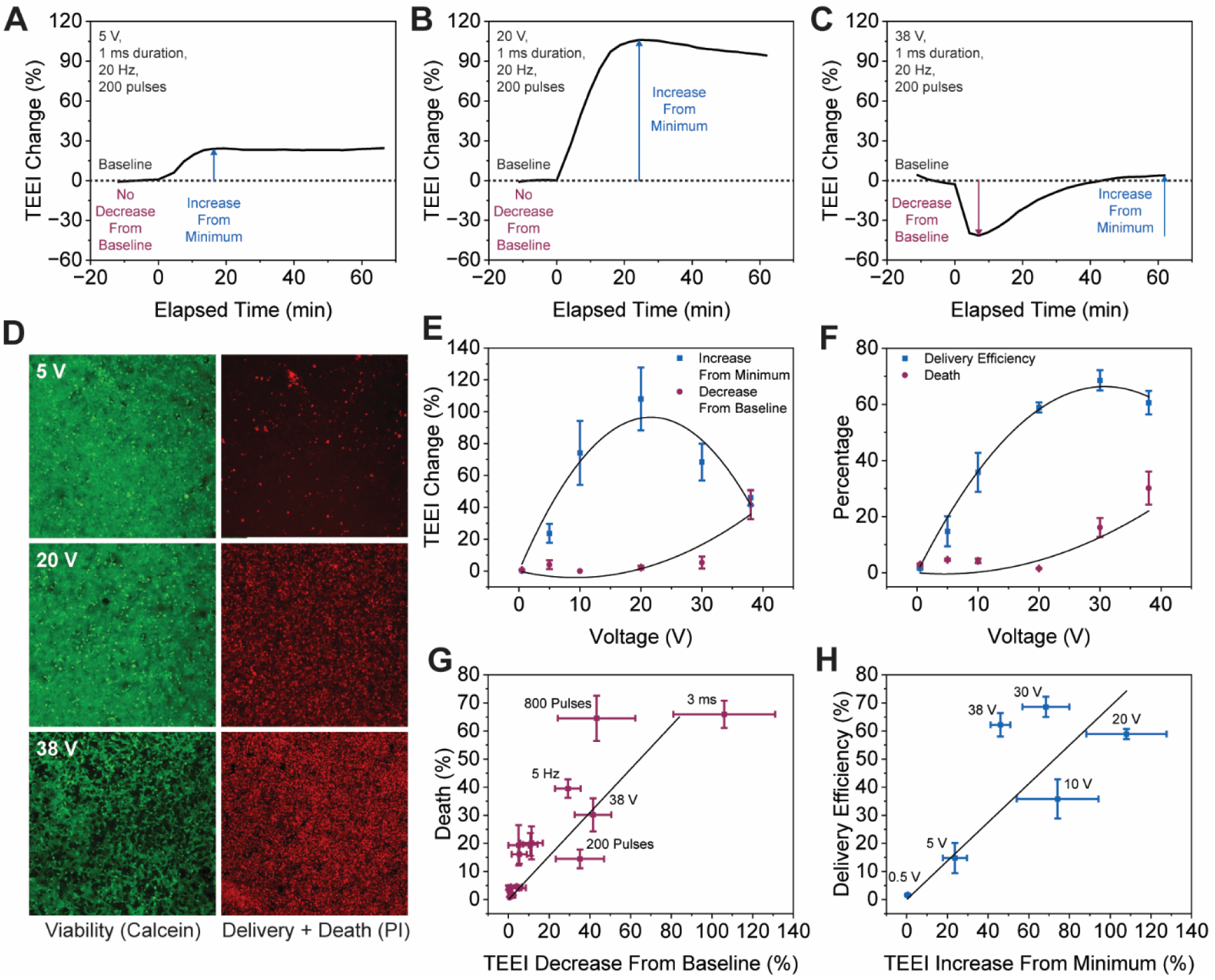
Components of the post-EP TEEI response are correlated to delivery metrics. **A.** Two TEEI response features labelled on the 5 V response. **B.** Two TEEI response features labelled on the 20 V response. **C.** Two TEEI response features labelled on the 38 V response. **D.** Fluorescent images showing viability and combined delivery and death at multiple voltages. **E.** TEEI increase from minimum and decrease at multiple voltages. **F.** Delivery efficiency and cell death at multiple voltages. **G.** Correlation between TEEI decrease and cell death for all data shown in Figure 3. Some conditions are unlabeled due to overlapping groups. **H.** Correlation between TEEI increase from minimum and PI delivery efficiency at multiple voltages. Error bars in D-F represent the SEM (n = 6).

To determine whether TEEI decrease was correlated with cell death, we plotted the results of all conditions tested in Figure 3 with their percent TEEI decrease from baseline on the x-axis and percent cell death (calculated as 100% minus viability) on the y-axis (Figure 4G). It should be noted that many of the conditions had almost no cell death or TEEI increase. There appears to be a positive correlation between TEEI decrease and cell death, which is significant because it means cell death can be estimated without fluorescent labels if the TEEI decrease is known. Having shown there is a correlation between cell death and TEEI decrease, we sought to determine whether there is also a correlation between delivery efficiency and TEEI increase. The increase from minimum is correlated with efficiency, but not delivery, because at higher voltages, smaller increases are seen but larger delivery occurs and viability decreases (Figure 4D). To determine whether the TEEI increase was correlated with delivery efficiency, we plotted delivery efficiency versus TEEI increase from minimum (Figure 4H). Delivery efficiency appears to be correlated with an increase in TEEI from minimum, therefore allowing delivery efficiency to be estimated using the TEEI response.

### Post-EP cell remodeling leads to an increase in transcellular impedance

To explain the post-EP TEEI response, we first considered which frequencies have the largest and smallest TEEI increase under our standard conditions. By comparing the TEEI at 100 Hz, 1 kHz, 10 kHz, and 100 kHz immediately before electroporation and at the maximum TEEI increase, we observed the 100 Hz TEEI increase was largest at 7.3 Ω·cm^2^ and was progressively smaller to 4.4 Ω·cm^2^ at 100 kHz (Figure 5A-B). Cell monolayer impedance is commonly represented using a resistor in parallel with a capacitor (*13, 14*). The resistor represents the paracellular impedance, the impedance experienced by current flowing around the cell through the cell-cell junctions (Figure 5C). The capacitor represents the transcellular impedance, the impedance experienced by current flowing through the cell membrane and cytoplasm (*13, 14*) (Figure 5C). Paracellular impedance typically dominates in the hundreds to thousands of Hz range, whereas transcellular impedance typically dominates in the tens of thousands of Hz (*14*). The largest TEEI increase that we observed is around 100 Hz, which suggests the increase may be primarily an increase in paracellular impedance.

**Figure 5.**
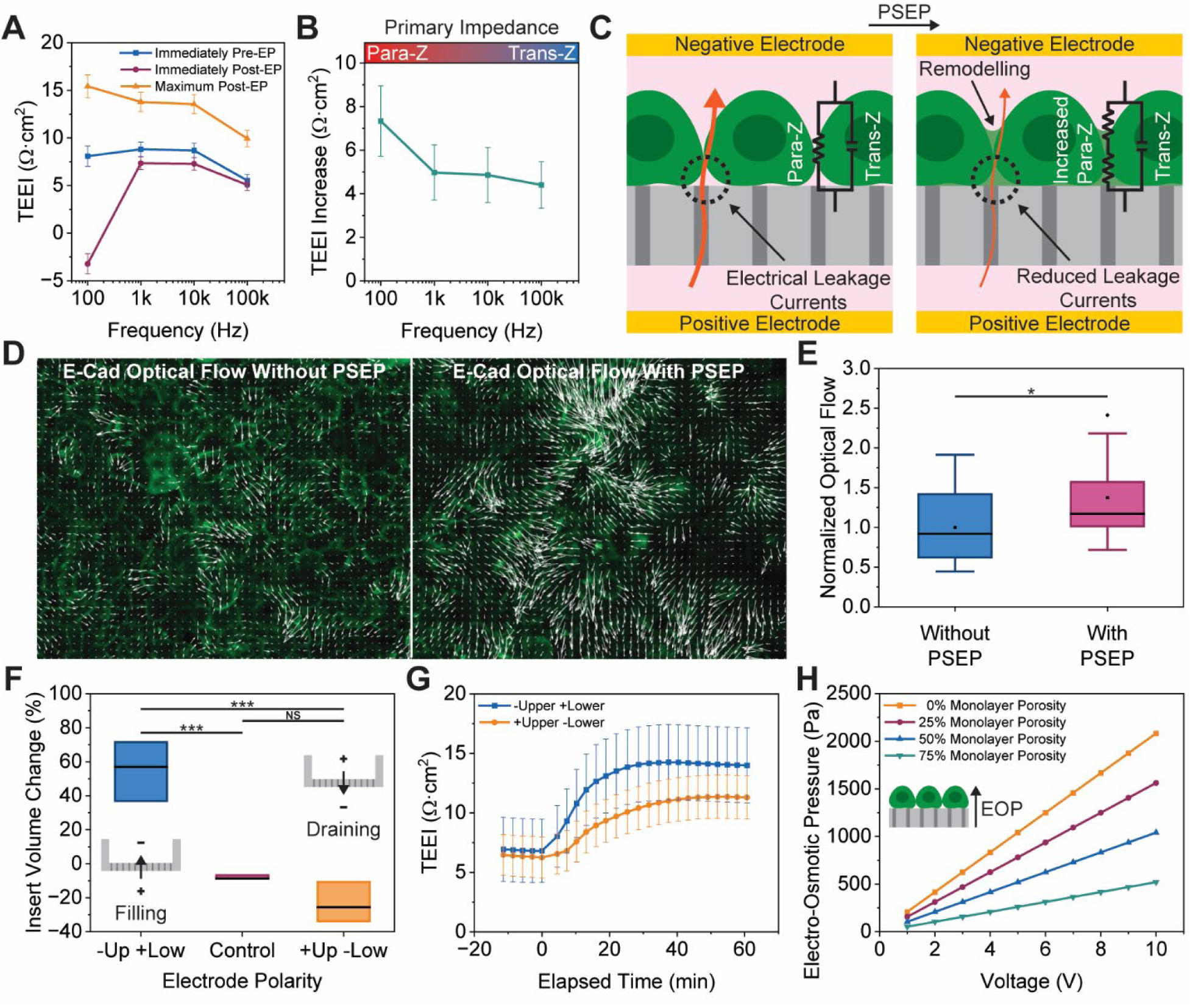
The post-EP TEEI response may be caused by cytoskeletal remodeling in response to electrical stimuli. **A.** TEEI at multiple frequencies before, after, and at the maximum post-EP time points. **B.** TEEI increase from pre-EP to maximum post-EP at multiple frequencies. **C.** Depiction of a cell monolayer before and after electroporation showing reduced leakage currents due to cytoskeletal remodeling and increased paracellular impedance (Para-Z) while transcellular impedance (Trans-Z) remains unchanged. **D.** Fluorescent images of GFP E-cadherin before and after PSEP with vector fields indicating optical flow. **E.** Comparison of normalized optical flow before and after PSEP. **F.** Change in insert volume from electro-osmotic transport of DI water without cells under different electrode polarities. Inset diagrams show electrode polarity relative to the insert and resulting electro-osmotic transport. **G.** Post-EP TEEI response with different electrode polarities. **H.** Simulation of electro-osmotic pressure applied to cell monolayer assuming different cell monolayer porosities. The inset diagram shows the direction of electro-osmotic pressure (EOP). Error bars in B (n = 21), C (n = 21), and H (n = 12) represent the SEM. For the box plots in F (n = 9) and G (n = 3), the shaded area represents the interquartile range (IQR), the error bars represent 1.5 times the IQR, and the horizontal line is the median.

We next investigated the cause of the increase in paracellular impedance by examining the cell remodeling after electroporation pulse. We hypothesize the confined electric field and fluid flow allow for more targeted stimuli and the adherent state allows cytoskeletal and cell-cell junction remodeling, which decreases leaky current and thus increases paracellular impedance (Figure 5C). Time lapses of A431 cells expressing green fluorescent protein (GFP) tagged E-cadherin were imaged before and after electroporation to determine whether electroporation resulted in significant morphological changes (Figure 5D, Movie S1 and S2). Although some cell movement and morphological changes are present prior to electroporation, following electroporation there is a statistically significant 37% average increase in cell monolayer movement due to the contraction and expansion of different regions (Figure 5E). This suggests PSEP may be stimulating the cell monolayer, inducing cell remodeling and increasing the cell monolayer impedance.

Having shown that cell junctions are affected by PSEP, we sought to investigate what stimulus may be responsible. As mentioned earlier, the electric field is one potential stimulus for cytoskeletal remodeling, but another potential stimulus is electro-osmotic flow. Electro-osmosis is bulk fluid flow caused by an electric field, in contrast with electrophoresis, which is related to the movement of charged molecules or particles within the electrolyte resulting from an electric field (*20*). Electrophoresis is often considered the dominant delivery mechanism during PSEP (*1–3, 9*), but the relative contributions to delivery of electrophoresis and electro-osmosis have not yet been demonstrated. To test whether electro-osmotic flow can occur in our substrates, we measured the amount of water transported through the substrate during the application of an electric field. To this end, the inserts were filled with DI water and weighed before and after pulsing to determine the change in water. It is worth mentioning that the lower ionic concentration in DI water compared with cell media increases the Debye length, which potentially favors electro-osmosis. This experiment revealed a 55.2% increase in insert volume under the standard negative upper electrode and positive lower electrode configuration, in contrast to a 23.3% decrease in volume when in reverse, and an 8.0% decrease in volume without pulses due to evaporation (Figure 5F). This water transport suggests electro-osmosis occurs in the system when a low ionic concentration is used, and it drives water transport towards the negative electrode.

We then sought to determine whether the lower transport with a positive upper electrode would translate to a lower post-EP TEEI response. Samples were electroporated with either the standard negative upper electrode and positive lower electrode configuration or the reversed configuration. Both conditions had similar pre-EP TEEI values as expected, but the reversed electrode polarity yielded a smaller response than our standard electrode polarity (Figure 5G). Next we evaluated how large of a mechanical stimulus electro-osmosis would exert on the cell monolayer if it occurs during PSEP. Our calculations suggest electro-osmosis may exert over 1 kPa of upward pressure on the cell monolayer (Figure 5H, Figure S5), which may be sufficient to induce a cell response (*21*).

## DISCUSSION

We have shown that TEEI measurements can be used without fluorescent imaging for label-free delivery. There are several advantages to label-free delivery including the ability to make pre- pulse adjustments, the ability to decouple delivery assessment from complications and restrictions imposed by cargo properties, improved temporal resolution for understanding transient behavior, and reduced cost and subjectivity compared to fluorescent imaging (*7*). By measuring impedance prior to electroporation, sample-to-sample variations in impedance due to factors such as confluency or cell type can be accounted for by adjusting waveform parameters. Label-free delivery also allows permeabilization to be measured in the absence of cargo, which removes confounding cargo parameters such as zeta potential, quantum yield, and toxicity resulting from cargo delivery rather than electrical exposure. When cargo delivery is desired, label-free delivery allows the use of unlabeled cargo or cargo at concentrations too low for imaging, which may be important for applications such as clinical research or industrial protein production. Furthermore, label-free delivery provides superior temporal resolution compared to fluorescent imaging because many PSEP systems are not capable of live imaging (*1, 9*), electrical measurements can be acquired faster than images, and electrical measurements do not require cargo to accumulate in sufficient concentrations prior to becoming visible (*7*). Label-free delivery also has the potential to cost less than imaging because impedance measurement equipment can be less expensive than fluorescent microscopes and initial waveform optimization can be performed without using expensive reagents (Figure S6). Despite these advantages, there are benefits to fluorescent imaging that TEEI cannot replicate due to the increased spatial information that imaging provides. PSEP analysis should ideally incorporate both TEEI and fluorescent imaging, with TEEI better suited to initial parametric optimization and understanding of temporal changes, and fluorescent imaging better for understanding final results.

The effect of electroporation on resistance measurements, and impedance measurements more generally, has been investigated in the literature. The application of electroporation pulses has been shown in many instances to cause a significant drop in cell impedance, followed by gradual recovery to baseline values, which is thought to be due to the permeabilization and resealing of the cell membrane (*10*). These impedance measurements have been performed before and after electroporation with diverse cell types *in vivo* (*22–26*) in tissues, and *in vitro* in monolayers (*27–34*), suspended pellets (*35*), and adherent (*36, 37*) and suspended (*38*) single-cells (see Table S5 for a more detailed summary). To our knowledge, this is the first study of impedance measurements before and after PSEP on non-silicon substrates, so the increase may have been observed in this study and not in others due to differences between PSEP on non-silicon substrates and other electroporation methods. The key difference between PSEP and most other electroporation methods is the existence of channels which confine the electric field and fluid flow. Furthermore, although there have been impedance studies of single-cell PSEP and other channel based methods such as nanostraw electroporation, these studies may not show the increase for two reasons: 1) silicon based substrates may be sufficiently different from track etched membranes to not experience the same response, particularly if the response is caused by electro-osmosis as a result of the track-etching process, and 2) if the response is caused by increases in paracellular impedance rather than transcellular impedance, it would only be evident in measurements of cell monolayers (more on this later).

To understand the differences between our observations and previous studies, it is also important to note the similarities. Both our observations and the literature show impedance increasing at a decreasing rate over similar time periods. The fundamental difference is that the literature consistently reports impedance rapidly decreasing before returning to baseline, whereas our observations show impedance increasing above baseline. Perhaps the simplest explanation of the discrepancy is that there are two distinct mechanisms causing the impedance increase (Figure 6). The gradual increase we are measuring may be recovery and resealing of the cell membrane as reported in other studies (*27, 31, 33, 34*), but the response is shifted upward by a second mechanism that occurs rapidly. The second mechanism may be occurring only during the application of the electric field, which is a relatively brief period (∼0.2%) of the total observation period. If we assume that the second mechanism occurs rapidly and shift our observed curves downward so that the final TEEI measurements are reduced proportionate to the reduction in viability, similar to what has been reported in other studies (*31*), it suggests the second mechanism is dependent on the voltage applied, as well as the viability or integrity of the cell monolayer.

**Figure 6.**
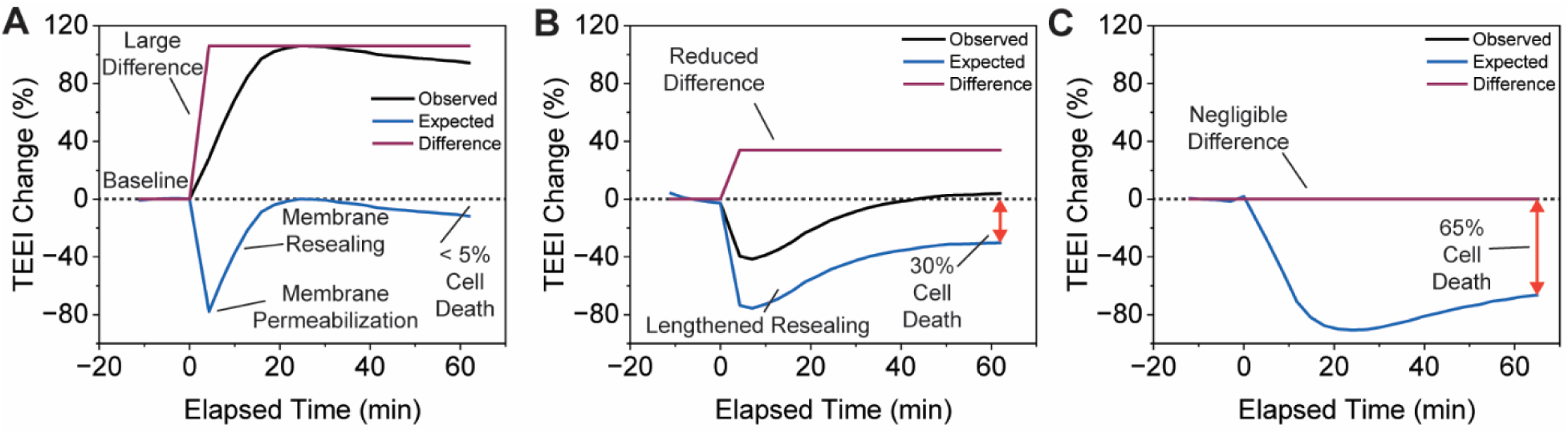
Post-EP TEEI response compared to the literature response. Depictions of the difference between our observations and responses similar to those reported in the literature. In each depiction, the literature curve is the observation curve shifted downward so that the TEEI does not exceed the initial TEEI and so the final TEEI is proportionate to the final viability. **A.** Depiction for waveforms that result in reversible electroporation of all cells. The observation curve is data from Figure 3Bi, 20 V. **B.** Depiction for waveforms that result in irreversible electroporation of some cells. The observation curve is data from Figure 3Bi, 38 V. **C.** Depiction for waveforms that result in irreversible electroporation of all cells. The observation curve is data from Figure 3Ei, 800 pulses. The observation curve is obscured by the overlapping expectation curve.

We postulate that the second mechanism could be the due to the focused electric field or electro-osmotic flow during PSEP serving as a mechanical stimulus, resulting in cytoskeletal remodeling that increases TEEI through an increase in paracellular impedance. Generally, paracellular impedance can increase from a reduction in paracellular space due to cell swelling (*39*) or tighter cell-cell junctions and cell-ECM adhesions (*14*). The porous substrate differentiates PSEP from other electroporation methods by allowing cells to remain adherent and by focusing the electric field and fluid flow through the channels. Specifically, the confined electric field and fluid flow may allow for more targeted stimuli, while the adherent state allows cytoskeletal and cell-cell junction remodeling as we have demonstrated in Figure 5D-E, which increases paracellular impedance. Moreover, this remodeling has been shown to increase TEEI (*12*), we observed HEK293T cells require a fibronectin coating for the post-EP TEEI increase (Figure S4), and electric fields have been shown to influence cell attachment, spreading, and contractility (*40–42*).

The idea of electro-osmotic flow as a stimulus is supported by studies showing electro-osmosis can occur within track-etched membranes (*43*). Electro-osmosis is thought to occur in track-etched membranes due to the negative surface charge on the channel walls from exposure of carboxyl groups during etching (*44*), as well as hydrophilic coatings applied during the manufacturing process such as polyvinylpyrrolidone (PVP). Positive ions accumulate along the negatively charged channel walls, forming a so-called Debye layer characterized by an excess of positive charge. When an electric field is applied along the channel, the positive ions within the Debye layer move towards the negative electrode, driving the flow within this layer and as a result, giving rise to the bulk flow (*20*). Such flow known as electro-osmosis flow is observed in our water transport experiment as shown in Figure 5F. We calculated that this electro-osmotic flow has the potential to exert sufficient mechanical stress, in the kPa range, to drive cell remodeling. Our calculations also show that the pressure induced by electro-osmosis is dependent on the porosity of the cell monolayer, which agrees with our data showing less confluent monolayers and those damaged by excessive electrical exposure resulted in smaller impedance increases. We have also shown that reversing the polarity of the electrode configuration results in a reversal of water transport and a reduced impedance change compared with the standard configuration, lending further support to electro-osmosis as a potential stimuli for cell remodeling.

We have also considered alternative explanations for the TEEI increase such as electro-osmotic compression (*45*) and ion channel synchronization (*46, 47*), however these explanations are unlikely because they rely on waveforms that are very different from the waveforms used in this study. We also considered that the TEEI increase could be due to electro-osmotic swelling, where electro-osmotic flow through the substrate enters the cell through the permeabilized membrane, causing an increase in cell volume, which results in a reduction of paracellular spaces and a corresponding increase in TEEI (*39*). However, even if an electro-osmotic flow is present, the small pores created in the cell membrane (*17*) may be insufficient to allow much flow into the cell. Indeed, we did not observe a significant increase in cell monolayer thickness before and after electroporation (Figure S7). The TEEI increase could also be from substrate channel blockage caused by the extraction of negatively charged intracellular biomolecules, however, channel blockage is also unlikely because the cytoplasm is primarily water so a large percentage of intracellular biomolecules would need to be extracted, which would likely result in much lower viability.

Some limitations of this study include our measurement device’s lack of phase detection and insufficient sampling rate to detect changes in impedance between individual electroporation pulses. Some researchers have utilized measurement devices with very high sample rates to show impedance changes not only after electroporation, but during electroporation (*32*). Phase detection is used with impedance measurement to separate impedance magnitude into real and imaginary components, which can be used with Nyquist plots for improved modelling of unknown impedances. Together, these improvements to our measurement device would provide more information about the nature of the post-EP response, adding clarity to the underlying response mechanism.

To summarize, we demonstrated an integrated TEEI-PSEP system and used it to record TEEI before and after electroporation. These measurements revealed a significant increase in TEEI following electroporation that contrasts with the temporary decrease in TEEI often reported in the literature. We comprehensively showed how different cell culture conditions and electroporation waveform parameters influence the post-EP TEEI response. Importantly, we demonstrated for the first time that features of the post-EP TEEI response can be used to estimate viability and delivery efficiency for label-free PSEP. Future study of the TEEI increase mechanism may provide further insight into PSEP and allow for greater optimization of delivery outcomes.

## MATERIALS AND METHODS

### Integrated impedance measurement and electroporation device design

Electrical signals for impedance measurement and electroporation were produced using a Keysight U2761A function generator. The output from the function generator was verified using a Tektronix TBS1062 oscilloscope and amplified using a Taidacent OPA541 amplifier (Figure S8). Root-mean-square voltage (V_rms_) was measured across known and unknown loads using a Keysight U2741A multimeter. A custom printed circuit board, or PCB, (PCBWay) was used as a control board to adjust which samples were being measured and pulsed by the multimeter and function generator (Figure S9). The control board contained Vishay VOR1121A6 solid-state relays, which were controlled using an Arduino Mega 2560 microcontroller. Logic gates were used on the control board to reduce the number of Arduino output pins needed. Matlab was used to control the function generator, multimeter, and Arduino, and to perform calculations and analysis of the impedance measurements. A Matlab graphical user interface (GUI) was created to control the system.

### Electrode array fabrication

The electrodes that formed the top and bottom of the electrode array consisted of custom PCBs (PCBWay) plated with 1 microinch thick electroless nickel immersion gold (ENIG) to ensure corrosion resistance and biocompatibility. Two Molex RJ25 surface mount modular connectors were soldered to the upper and lower PCBs for connection to the control board. 3D printed covers were placed over the pin electrodes to ensure measurements were not influenced by fluctuations in cell culture media volume. The pin covers and wells were printed on a Formlabs Form 3B 3D printer using BioMed Clear Resin. Biocompatible Viton fluorelastomer O-rings were placed within grooves on the bottom of the wells and the wells were fastened to the lower PCBs. The wells remained fastened to the lower PCBs for 20 uses, at which point the PCBs were replaced due to corrosion (Table S6).

### Cell culture

Greiner Bio-One 24-well inserts with polyethylene terephthalate (PET) substrates containing 400 nm diameter pores at a density of 2·10^6^ pores·cm^-2^ were coated with 100 µL of 1 µg/mL human plasma fibronectin (Sigma-Aldrich) in PBS (Gibco) and incubated for 3 hours at 37 °C. Coated substrates were washed twice with distilled water and once with cell culture media prior to seeding to remove excess fibronectin. Cells were seeded at least 12 hours prior to electroporation to allow sufficient adherence. A431 cells were seeded at 200,000 cells·cm^-2^ and HEK293T cells were seeded at 400,000 cells·cm^-2^. Cell culture media for both A431 and HEK293T cells consisted of Dulbecco’s modified Eagle medium (DMEM) (Gibco) with 10% v/v fetal bovine serum (FBS) (Gibco) and 1% v/v penicillin-streptomycin (Gibco).

### Impedance measurement

An alternating current (AC) sine wave was applied across a resistor and an unknown load in series, and the V_rms_ across the unknown load was measured. The device used relays on the control board to cycle through increasingly larger resistors until the V_rms_ across the unknown load was closest to half the total input V_rms_. The V_rms_ was then measured across the chosen resistor and used in conjunction with the previously measured unknown load V_rms_ using the following equation, where |Z_unknown_| is the impedance magnitude of the unknown load, V_unknown_ is the V_rms_ across the unknown load, V_R_ is the V_rms_ across the chosen resistor, and |Z_R_| is the impedance magnitude of the resistor.

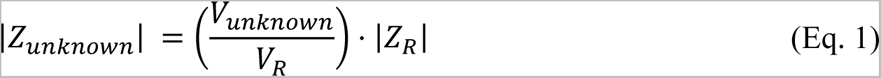

The resistors were 100 Ω, 1 kΩ, 10 kΩ, 50 kΩ, and 100 kΩ, and measurement would begin using the lowest resistor. 0.5 V was chosen for the measurement amplitude because it was high enough to achieve a large signal to noise ratio without being high enough to cause damage to the cells.

### Device validation using a TEER meter and an LCR meter

To ensure our device could accurately measure TEER, our device was compared to a World Precision Instruments (WPI) Epithelial Volt/Ohm Meter 3 (EVOM3) TEER meter. Both devices were connected to a WPI EndOhm chamber to provide a common electrode interface for comparison. A431 cells were seeded in the inserts at 200,000 cells·cm^-2^ for over 12 hours. The EndOhm chamber contained 1 mL of cell culture media and the inserts each contained 200µL of cell culture media. Each device measured across inserts with and without cells. The same inserts were measured with both devices. Our device measured the impedance at 12.5 Hz, which is the same frequency the EVOM3 measures at. To verify impedance measurement accuracy across multiple frequencies, our device was compared to a Fluke PM6306 LCR meter. Multiple configurations of resistors and capacitors were measured with both devices at 50 Hz, 100 Hz, 500 Hz, 1 kHz, 2.5 kHz, 5 kHz, 10 kHz, 25 kHz, 50 kHz, and 100 kHz.

### Measurement of change in TEEI following electroporation

Electrode arrays were sterilized using 70% ethanol for 10 minutes prior to each use. Inserts were placed into each well in the electrode array and the upper PCB was fastened to the wells. Small holes in the upper PCB over each well allowed for gas exchange during the experiments. The electrode array was placed in an incubator and connected to cables from the measurement and electroporation device. The electrode array was allowed to thermally equilibrate in the incubator for 1 hour to ensure there were no impedance decreases due to an increase in temperature. Inserts were typically placed in multiple electrode arrays at a time to allow the next electrode array to equilibrate during the measurements and electroporation of the previous electrode array. After 1 hour, the experimental parameters were entered into the GUI and the measurements and electroporation were started. Parameters that could be adjusted with the GUI included measurement frequencies, measurement voltage, pulse voltage, pulse duration, pulse frequency, pulse number, train number, train frequency, and the duration of measurements before and after pulsing. The standard protocol involved 10 minutes of impedance measurements at 100 Hz, 1 kHz, 10 kHz, and 100 kHz, followed by electroporation with unipolar square waves at 30 V, 1 ms pulse duration, 20 Hz, and 200 pulses, followed by an additional 60 minutes of impedance measurements at the previous 4 frequencies. The measurement rate was one measurement every 6.8 seconds, or 147 mHz. TEEI values were calculated using the following equation, where A is the substrate area (0.336 cm^2^ for the inserts used in this study), Z_cells_ is the impedance magnitude of inserts containing cells, and Z_nocells_ is the impedance magnitude of inserts without cells.

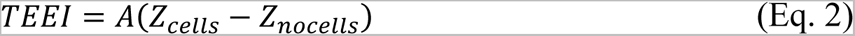

### Viability and delivery quantification

To assess whether electroporation was occurring, we added PI to the wells before applying various waveforms. PI is typically used in conjunction with calcein in a live-dead assay because calcein relies on enzymatic cleavage to label living cells while PI is membrane impermeable and therefore only labels dead cells. However, when PI is added extracellularly prior to electroporation, simultaneous labelling with both calcein and PI is indicative of delivery caused by electroporation. Cells were stained with 12.3 µg/mL (20 µM) Hoechst 33342 (Thermo Scientific), 2.5 µg/mL (2.5 µM) calcein AM (Invitrogen), and 5 µg/mL (7.5 µM) PI (Invitrogen) and incubated for 5 minutes prior to imaging with a Zeiss Axio Observer 5 fluorescent microscope. Except for experiments evaluating PI delivery during electroporation, PI was added 1 hour after electroporation to ensure internalization was due to cell death, not delivery. For experiments evaluating PI delivery, 0.1 mg/mL of PI was added to cell culture media in the cargo chamber prior to electroporation. Image processing was performed using a custom CellProfiler pipeline. Hoechst 33342 was used for quantification of total cell count, calcein was used for quantification of confluency, and PI was used for quantification of viability. The total number of cells was determined by counting the number of cells labelled with Hoechst 33342. Viability was calculated as the number of cells not labelled with PI divided by the total number of cells, and confluency was calculated as the area of the image stained with calcein divided by the total image area. For experiments measuring PI delivery, viability was calculated as the number of cells labelled with calcein divided by the total number of cells, delivery was calculated as the number of cells labelled with PI divided by the total number of cells, and delivery efficiency was calculated as the number of cells labelled with both calcein and PI divided by the total number of cells.

### Measurement of optical flow before and after electroporation

A431 cells expressing GFP E-cadherin were cultured on inserts using the standard conditions. A 10-minute time lapse was recorded for each insert prior to electroporation with one image taken each minute. The inserts were then individually electroporated using 20V while keeping the other waveform parameters standard. A second 10-minute time lapse was recorded for each insert immediately following electroporation. A custom Matlab code was used to measure the optical flow in each time lapse and the tical flow of both the pre- and post-EP time lapses were normalized to the average of the pre-EP time lapses.

### Electro-osmotic transport of deionized water

Inserts were weighed before and after adding 150 µg of deionized water. 1 mL of deionized water was added to the well. 500 pulses of 400 V, 100 ms duration, and 5 Hz were applied to the insert in the well using a Bio-Rad Gene Pulser II with RF Module. After applying the pulses, the insert with water was weighed to determine the final volume.

### Measurement of cell monolayer thickness

Inserts seeded and cultured with cells under the standard cell culture conditions were divided into 2 groups: a negative control group that would not be electroporated, and a group that would be electroporated with the standard waveforms and undergo the standard post-EP impedance measurement. Afterwards, cells were stained with 2.5 µg/mL (2.5 µM) calcein AM and incubated for 5 minutes. Z-stacks were imaged using a Zeiss LSM 800 laser scanning confocal microscope. Matlab was used to determine the median intensity of each image in each Z stack. The median intensities for each Z-stack were then normalized to the brightest image in each Z-stack. A cutoff of 25% of the maximum median intensity was used to determine the upper and lower boundaries of the cell monolayer. Cell monolayer height was determined using the number of images in each monolayer and the known spacing between each image.

### Statistics

Statistical analysis was performed using Origin. For the comparison between the TEER meter measurements and the TEEI-PSEP measurements, a two-sample t-test was used. For all other figures, normality and equality of group variances were assessed prior to calculation of one-way analysis of variance (ANOVA) with Bonferroni pairwise comparisons. Normality was assessed using the Shapiro-Wilk test. Equality of group variances was assessed using the Brown-Forsythe test. For instances of non-normality, the Kruskal-Wallis test was used with Dunn’s test for pairwise comparisons. For instances of unequal variances, Welch’s ANOVA was used with Games-Howell pairwise comparisons.

## ACKNOWLEDGEMENTS

We acknowledge the funding support from the NSF (Awards 1826135, 1936065, 2143997), the NIH National Institutes of General Medical Sciences P20GM113126 (Nebraska Center for Integrated Biomolecular Communication) and P30GM127200 (Nebraska Center for Nanomedicine), the Nebraska Collaborative Initiative and the Voelte-Keegan Bioengineering Support. Manufacturing of the device was performed at the NanoEngineering Research Core Facility (NERCF), which is partially funded by the Nebraska Research Initiative.

## AUTHOR CONTRIBUTIONS

J.R.B., T.C.H., S.L., S.M., J.S.P., R.Y. conceived and designed the analysis, J.R.B., T.C.H., S.R.L., I.M. collected the data, J.R.B., T.C.H., S.R.L., S.M., J.S.P., R.Y. performed the analysis, J.R.B., T.C.H., S.M., J.S.P., R.Y. wrote the paper.

## DATA AVAILABILITY

All data needed to evaluate the conclusions in the paper are present in the paper and/or the Supplementary Materials. Additional data and materials are available from the corresponding author upon request

## References

1. P. Mukherjee, C.-Y. Peng, T. McGuire, J. W. Hwang, C. H. Puritz, N. Pathak, C. A. Patino, R. Braun, J. A. Kessler, H. D. Espinosa, Single cell transcriptomics reveals reduced stress response in stem cells manipulated using localized electric fields. Materials Today Bio 19, 100601 (2023).

2. N. Pathak, C. A. Patino, N. Ramani, P. Mukherjee, D. Samanta, S. B. Ebrahimi, C. A. Mirkin, H. D. Espinosa, Cellular Delivery of Large Functional Proteins and Protein– Nucleic Acid Constructs via Localized Electroporation. Nano Letters 23, 3653–3660 (2023).

3. C. A. Patino, P. Mukherjee, E. J. Berns, E. H. Moully, L. Stan, M. Mrksich, H. D. Espinosa, High-Throughput Microfluidics Platform for Intracellular Delivery and Sampling of Biomolecules from Live Cells. ACS Nano 16, 7937–7946 (2022).

4. J. Brooks, G. Minnick, P. Mukherjee, A. Jaberi, L. Chang, H. D. Espinosa, R. Yang, High Throughput and Highly Controllable Methods for In Vitro Intracellular Delivery. Small 16, 2004917 (2020).

5. Z. Dong, S. Yan, B. Liu, Y. Hao, L. Lin, T. Chang, H. Sun, Y. Wang, H. Li, H. Wu, Single Living Cell Analysis Nanoplatform for High-Throughput Interrogation of Gene Mutation and Cellular Behavior. Nano Letters, (2021).

6. Y. Cao, E. Ma, S. Cestellos-Blanco, B. Zhang, R. Qiu, Y. Su, J. A. Doudna, P. Yang, Nontoxic nanopore electroporation for effective intracellular delivery of biological macromolecules. Proceedings of the National Academy of Sciences 116, 7899 (2019).

7. Y. Ye, X. Luan, L. Zhang, W. Zhao, J. Cheng, M. Li, Y. Zhao, C. Huang, Single-Cell Electroporation with Real-Time Impedance Assessment Using a Constriction Microchannel. Micromachines 11, 10.3390/mi11090856 (2020).

8. C. A. Patino, N. Pathak, P. Mukherjee, S. H. Park, G. Bao, H. D. Espinosa, Multiplexed high-throughput localized electroporation workflow with deep learning–based analysis for cell engineering. Science Advances 8, eabn7637.

9. J. R. Brooks, I. Mungloo, S. Mirfendereski, J. P. Quint, D. Paul, A. Jaberi, J. S. Park, R. Yang, An equivalent circuit model for localized electroporation on porous substrates. Biosensors and Bioelectronics 199, 113862 (2022).

10. Q. Castellví, B. Mercadal, A. Ivorra, "Assessment of Electroporation by Electrical Impedance Methods" in Handbook of Electroporation, D. Miklavčič, Ed. (Springer International Publishing, Cham, 2017), pp. 671–690.

11. S.-E. Choi, H. Khoo, S. C. Hur, Recent Advances in Microscale Electroporation. Chemical Reviews 122, 11247–11286 (2022).

12. L. Cacopardo, J. Costa, S. Giusti, L. Buoncompagni, S. Meucci, A. Corti, G. Mattei, A. Ahluwalia, Real-time cellular impedance monitoring and imaging of biological barriers in a dual-flow membrane bioreactor. Biosensors and Bioelectronics 140, 111340 (2019).

13. B. Srinivasan, A. R. Kolli, M. B. Esch, H. E. Abaci, M. L. Shuler, J. J. Hickman, TEER Measurement Techniques for In Vitro Barrier Model Systems. Journal of Laboratory Automation 20, 107–126 (2015).

14. K. Benson, S. Cramer, H.-J. Galla, Impedance-based cell monitoring: barrier properties and beyond. Fluids and Barriers of the CNS 10, 5 (2013).

15. J. P. Vigh, A. Kincses, B. Ozgür, F. R. Walter, A. R. Santa-Maria, S. Valkai, M. Vastag, W. Neuhaus, B. Brodin, A. Dér, M. A. Deli, Transendothelial Electrical Resistance Measurement across the Blood–Brain Barrier: A Critical Review of Methods. Micromachines 12, 10.3390/mi12060685 (2021).

16. T. Vindiš, A. Blažič, D. Khayyat, T. Potočnik, S. Sachdev, L. Rems, Gene Electrotransfer into Mammalian Cells Using Commercial Cell Culture Inserts with Porous Substrate. Pharmaceutics 14, 10.3390/pharmaceutics14091959 (2022).

17. P. Mukherjee, S. S. P. Nathamgari, J. A. Kessler, H. D. Espinosa, Combined Numerical and Experimental Investigation of Localized Electroporation-Based Cell Transfection and Sampling. ACS Nano 12, 12118–12128 (2018).

18. Z. Fei, Y. Wu, S. Sharma, D. Gallego-Perez, N. Higuita-Castro, D. Hansford, J. J. Lannutti, L. J. Lee, Gene Delivery to Cultured Embryonic Stem Cells Using Nanofiber-Based Sandwich Electroporation. Analytical Chemistry 85, 1401–1407 (2013).

19. P. Mukherjee, E. J. Berns, C. A. Patino, E. Hakim Moully, L. Chang, S. S. P. Nathamgari, J. A. Kessler, M. Mrksich, H. D. Espinosa, Temporal Sampling of Enzymes from Live Cells by Localized Electroporation and Quantification of Activity by SAMDI Mass Spectrometry. Small 16, e2000584 (2020).

20. J. L. Anderson, Colloid Transport by Interfacial Forces. Annual Review of Fluid Mechanics 21, 61–99 (1989).

21. T. Iskratsch, H. Wolfenson, M. P. Sheetz, Appreciating force and shape-the rise of mechanotransduction in cell biology. Nat Rev Mol Cell Biol 15, 825–833 (2014).

22. U. Pliquett, R. Langer, J. C. Weaver, Changes in the passive electrical properties of human stratum corneum due to electroporation. Biochimica et Biophysica Acta (BBA) - Biomembranes 1239, 111–121 (1995).

23. D. Cukjati, D. Batiuskaite, F. André, D. Miklavčič, L. M. Mir, Real time electroporation control for accurate and safe in vivo non-viral gene therapy. Bioelectrochemistry 70, 501–507 (2007).

24. A. Ivorra, B. Rubinsky, In vivo electrical impedance measurements during and after electroporation of rat liver. Bioelectrochemistry 70, 287–295 (2007).

25. Y. Granot, B. Rubinsky, Methods of optimization of electrical impedance tomography for imaging tissue electroporation. Physiological Measurement 28, 1135 (2007).

26. A. Ivorra, B. Al-Sakere, B. Rubinsky, L. M. Mir, In vivo electrical conductivity measurements during and after tumor electroporation: conductivity changes reflect the treatment outcome. Physics in Medicine & Biology 54, 5949 (2009).

27. J. A. Stolwijk, C. Hartmann, P. Balani, S. Albermann, C. R. Keese, I. Giaever, J. Wegener, Impedance analysis of adherent cells after in situ electroporation: Non-invasive monitoring during intracellular manipulations. Biosensors and Bioelectronics 26, 4720–4727 (2011).

28. Z. J. Xing, D. G. Rong, Z. Jie, Z. Y. Chun, Z. Y. Jun, G. G. Zhen, Detrimental Effect of Electromagnetic Pulse Exposure on Permeability of In Vitro Blood-brain-barrier Model. Biomedical and Environmental Sciences 26, 128 (2013).

29. T. García-Sánchez, M. Guitart, J. Rosell-Ferrer, A. M. Gómez-Foix, R. Bragós, A new spiral microelectrode assembly for electroporation and impedance measurements of adherent cell monolayers. Biomedical Microdevices 16, 575–590 (2014).

30. T. García-Sánchez, A. Azan, I. Leray, J. Rosell-Ferrer, R. Bragós, L. M. Mir, Interpulse multifrequency electrical impedance measurements during electroporation of adherent differentiated myotubes. Bioelectrochemistry 105, 123–135 (2015).

31. X. Guo, R. Zhu, Controllable in-situ cell electroporation with cell positioning and impedance monitoring using micro electrode array. Scientific Reports 6, 31392 (2016).

32. T. García-Sánchez, R. Bragós, L. M. Mir, In vitro analysis of various cell lines responses to electroporative electric pulses by means of electrical impedance spectroscopy. Biosensors and Bioelectronics 117, 207–216 (2018).

33. J. A. Stolwijk, J. Wegener, Impedance analysis of adherent cells after in situ electroporation-mediated delivery of bioactive proteins, DNA and nanoparticles in µL- volumes. Scientific Reports 10, 21331 (2020).

34. D. Lee, S. S. Y. Chan, N. Aksic, N. Bajalovic, D. K. Loke, Ultralong-Time Recovery and Low-Voltage Electroporation for Biological Cell Monitoring Enabled by a Microsized Multipulse Framework. ACS Omega 6, 35325–35333 (2021).

35. I. G. Abidor, L. H. Li, S. W. Hui, Studies of cell pellets: II. Osmotic properties, electroporation, and related phenomena: membrane interactions. Biophysical Journal 67, 427–435 (1994).

36. A. G. Pakhomov, J. F. Kolb, J. A. White, R. P. Joshi, S. Xiao, K. H. Schoenbach, Long-lasting plasma membrane permeabilization in mammalian cells by nanosecond pulsed electric field (nsPEF). Bioelectromagnetics 28, 655–663 (2007).

37. D. Xu, J. Fang, H. Wang, X. Wei, J. Yang, H. Li, T. Yang, Y. Li, C. Liu, N. Hu, Scalable Nanotrap Matrix Enhanced Electroporation for Intracellular Recording of Action Potential. Nano Letters 22, 7467–7476 (2022).

38. S. C. Bürgel, C. Escobedo, N. Haandbæk, A. Hierlemann, On-chip electroporation and impedance spectroscopy of single-cells. Sensors and Actuators B: Chemical 210, 82–90 (2015).

39. H. Albus, J. A. Groot, J. Siegenbeek van Heukelom, Effects of glucose and ouabain on transepithelial electrical resistance and cell volume in stripped and unstripped goldfish intestine. Pflügers Archiv 383, 55–66 (1979).

40. K. Katoh, Effects of Electrical Stimulation on the Signal Transduction-Related Proteins, c-Src and Focal Adhesion Kinase, in Fibroblasts. Life 12, 10.3390/life12040531 (2022).

41. S. Staehlke, M. Bielfeldt, J. Zimmermann, M. Gruening, I. Barke, T. Freitag, S. Speller, U. Van Rienen, B. Nebe, Pulsed Electrical Stimulation Affects Osteoblast Adhesion and Calcium Ion Signaling. Cells 11, 10.3390/cells11172650 (2022).

42. P. M. Graybill, A. Jana, R. K. Kapania, A. S. Nain, R. V. Davalos, Single Cell Forces after Electroporation. ACS Nano 15, 2554–2568 (2021).

43. P. J. Kemery, J. K. Steehler, P. W. Bohn, Electric Field Mediated Transport in Nanometer Diameter Channels. Langmuir 14, 2884–2889 (1998).

44. C. Yang, "Ion and Particle Transport in Nanopores", thesis, University of California, Irvine (2018).

45. T. B. Saw, X. Gao, M. Li, J. He, A. P. Le, S. Marsh, K.-h. Lin, A. Ludwig, J. Prost, C. T. Lim, Transepithelial potential difference governs epithelial homeostasis by electromechanics. Nature Physics 18, 1122–1128 (2022).

46. V. Tran, X. Zhang, L. Cao, H. Li, B. Lee, M. So, Y. Sun, W. Chen, M. Zhao, Synchronization Modulation Increases Transepithelial Potentials in MDCK Monolayers through Na/K Pumps. PLOS ONE 8, e61509 (2013).

47. P. Liang, J. Mast, W. Chen, Synchronization Modulation of Na/K Pumps Induced Membrane Potential Hyperpolarization in Both Physiological and Hyperkalemic Conditions. The Journal of Membrane Biology 252, 577–586 (2019).

