## Supplemental Information for "Transepithelial Electrical Impedance Increase Following Porous Substrate Electroporation Enables Label-Free Delivery"

Justin R. Brooks *et al.*

*Corresponding author:
Ruiguo Yang,

**This PDF file includes:**

Supplementary Text
Figs. S1 to S9
Tables S1 to S6
Movies S1 to S2
References (1 to 23)

**Other Supplementary Materials for this manuscript include the following:**

Movies S1 to S2

### SUPPLEMENTAL TEXT

#### Device validation

Prior to measuring TEEI following electroporation, we designed a system capable of both measuring TEEI and performing PSEP (Figure 1 in the main text). There are commercially available TEEI systems such as nanoAnalytics' cellZscope and Applied Biosystems' ECIS Z-Theta, but to our knowledge they cannot produce electrical waveforms used in PSEP. Moreover, commercial TEEI electrodes are not designed to withstand the corrosion that occurs during electroporation, and routinely replacing the electrodes would be prohibitively expensive. Developing our own device also provided much more customization than working with a commercial system.

We previously demonstrated a voltage divider can be used to measure the PSEP system impedance (*1*). Due to their simplicity, voltage dividers can be used with a variety of impedances and waveforms. However, voltage dividers are inaccurate when the known load and unknown load do not have similar impedances. Wheatstone bridges have been used as an alternative due to their improved accuracy but can be more complicated to use when the applied waveform or the behavior of the unknown load have yet to be determined. Instead, our system utilizes a variable load voltage divider where the known load is automatically adjusted to have a similar impedance to the unknown load, ensuring consistently high accuracy.

To determine how accurately our device can measure TEER, we compared our device to the World Precision Instruments Epithelial Voltohmmeter 3 (EVOM3). (Figure S1C). The difference between the measurements made by our device and the EVOM3 is remarkably small (1.9%), indicating that our device can reliably measure TEER values. The TEER measured by both devices is smaller than TEER values typically reported in the literature, but TEER values of similar magnitude have been reported (*2*). Experimental conditions such as chamber electrodes, 24-well inserts, the cell line used, and the shorter culture period of 12 to 24 hours were chosen to reflect common PSEP conditions but may contribute to the lower TEER values (*3, 4*). To assess whether our device could accurately measure impedances across a range of frequencies relevant to PSEP, we compared our device to a Fluke PM6306 LCR meter (Figure S1D-F, Table S1). Various configurations of resistors and capacitors were measured at frequencies ranging from 50 Hz to 100 kHz. The measurements from our device were in close agreement with the measurements from the LCR meter, with an average difference of 0.7%, showing our device can accurately measure impedance across a range of frequencies.

#### Effects of non-cellular changes on impedance increase

After observing the TEEI increase, we sought to determine whether the increase could be attributed to a noncellular explanation such as temperature change, corrosion, or bubble formation. First, we considered temperature change, but the impedance of a solution is inversely correlated with temperature, so an impedance increase requires a temperature decrease. However, the only likely change in temperature following electroporation would be a slight temperature *increase* due to joule heating (*5*). Furthermore, the effect only occurs with cells present, which would not affect joule heating or any known cooling mechanism, and if the increase was due to a temperature decrease, the impedance should decay similarly to the impedance decay that occurs when the samples are first placed in the incubator from room temperature (Figure S3A).

We then tested whether electrode corrosion during pulsing could be causing the impedance increase. An electrode array was incubated with cell culture media and inserts without cells. The system impedance was measured before applying the electroporation waveforms, then measured again. This process was repeated 100 times to observe how the waveforms corroded the electrodes and increased the impedance (Figure S3B-C, Table S6). Although the impedance increased, the increase was too small to explain the response and we accounted for this response in all TEEI data by using negative control groups in each experiment. Any increase due to corrosion would occur in both the experimental groups and the negative control groups and the increase would be negated when the impedance of the negative control groups is subtracted from the impedance of the experimental groups.

We also considered whether bubbles could be causing the impedance increase, but bubbles were not observed following electroporation. Considering the impedance increased only with cells present, we evaluated whether the cell monolayer could be passively increasing impedance by serving as a barrier between the two electrodes. For example, perhaps the impedance increase is due to the formation of microbubbles on the lower electrode that float into the substrate channels and become trapped by the cell monolayer. To test this hypothesis, we fixed cells with 4% paraformaldehyde for 10 minutes prior to electroporation to observe whether the impedance would still increase (Figure S3D). Following fixation, the cells were washed twice with PBS and incubated for 1 hour in cell culture media as usual. The impedance started at a higher initial value but did not increase following electroporation, suggesting the impedance increase is due to an active response in the cell monolayer.

**Calculation of electro-osmosis induced mechanical stress**

We sought to investigate the electro-osmosis as a potential underlying mechanism for the increase in TEEI response (Figure 5 in the main text). Electro-osmotic flow is a bulk fluid motion driven by an external electric field in charged channels. The external electric field exerts the force on excess counter-ions in a diffusive Debye layer of the charged surface, consequently driving the motion of ions that leads to the electro-osmotic flow within the bulk fluid. Under the assumption of a thin Debye layer thickness $\lambda_D$ with respect to the characteristic length scale of the channel, the effect of such a force within the Debye layer can be modeled as an effective slip velocity at the boundaries. This assumption is particularly relevant to the current case of study, given that the pore diameter is significantly larger than the Debye layer thickness (i.e., $H = 400\, nm \gg \lambda_D = 0.7\, nm - 8\, nm$). With this assumption, the electro–osmosis velocity profile takes a plug–like form and can be expressed as:

$$u^{eo} = \frac{\varepsilon \zeta_w}{\eta} E_t$$

where $\eta$ is the viscosity of the electrolyte, and $E_t$ is the local tangential electric field. The parameter $\zeta_w$ is the channel wall zeta potential, which can be interpreted as the potential drop across the Debye layer (*6*). Using the current experimental parameters, the electro-osmosis velocity $u^{eo}$ is calculated and plotted in Figure S5A as a function of the voltage difference across the channel length. With this electro-osmosis velocity, we can calculate the total flow rate across the membrane, given the membrane area of $0.336\ cm^2$ and a channel density of 2e6 channels per $cm^2$. Figure S5B shows the time-averaged flow rate $Q^{eo}$, considering a 2% duty cycle, as a function of the voltage difference. Both the velocity and the flow rate exhibit a linear increase with the voltage difference. It is important to note that this relationship is particularly intuitive when considering that the predominant driving mechanism is the linear electrokinetic phenomena.

The presence of the cell monolayer, attached to the membrane at the flow outlet, can partially or fully obstruct the electro-osmotic flow, depending on the porosity of the cell monolayer. This flow blockage is likely to give rise to back pressure and mechanical stress at the cell monolayer. We refer to this induced back pressure as an "electro-osmosis pressure," denoted as $\Delta P$. This electro-osmosis pressure primarily depends on the electro-osmosis velocity and the porosity of the cell monolayer ($\epsilon$). Using the following relation,

$$\frac{\Delta P \pi d^4}{128 \eta L} = (1-\epsilon) Q^{eo}$$

where $d$ is the pore diameter and $L$ is the channel length. the electro-osmosis pressure can be obtained by

$$\Delta P = 32 \frac{\eta L}{d^2} u^{eo} (1-\epsilon)$$

With this expression, we have plotted the electro-osmosis pressure as a function of the voltage difference for different monolayer porosities in Figure 5H in the main text.

### SUPPLEMENTAL FIGURES

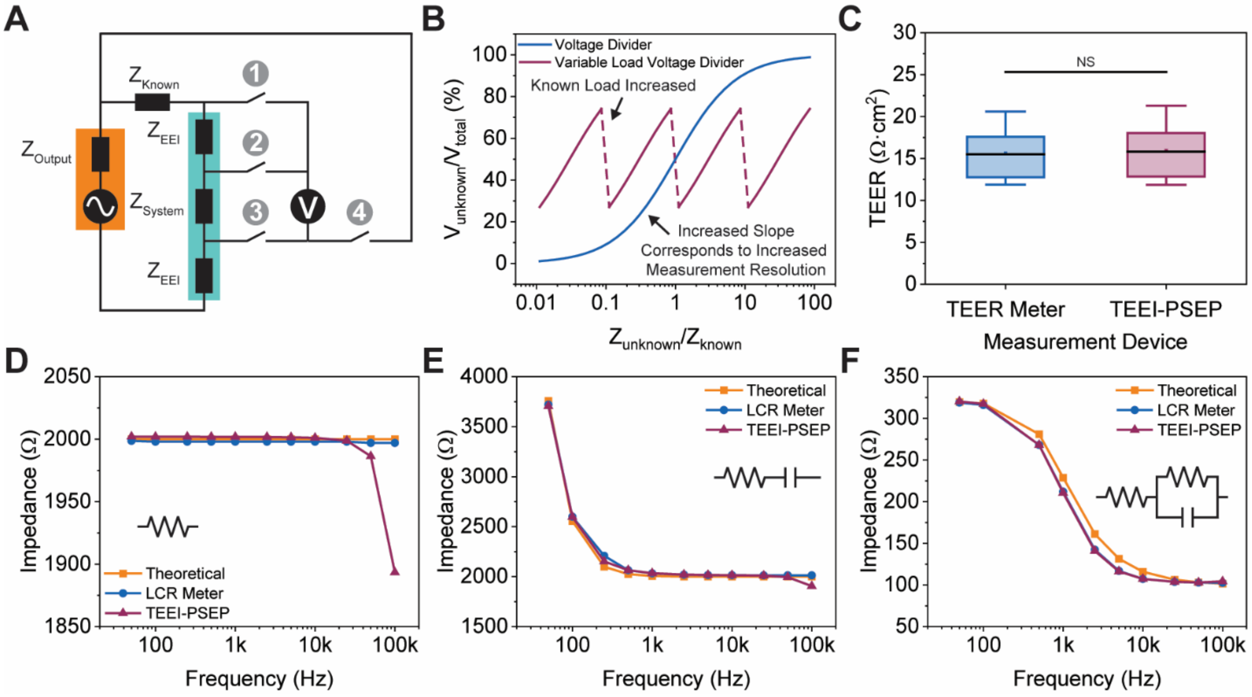


**Figure S1. Measurement System Design and Validation.** Overview of the automated voltage divider measurement system and comparisons to commercial measurement systems. **A.** Simplified circuit diagram of the system illustrating how voltage divider measurements are performed. The function generator is shown in orange, and the electrochemical cell is shown in green. Switches 1 and 4 are closed during measurement across the known load, and switches 2 and 3 are closed during 4 electrode measurement of the PSEP system. **B.** Theoretical comparison of a standard voltage divider and the variable load voltage divider used in our system. **C.** Comparison between TEER measurements in an EndOhm chamber using an EVOM3 TEER meter and our TEEI-PSEP system. The shaded areas represent the interquartile ranges (IQRs), the error bars represent 1.5 times the IQRs, the horizontal lines are the medians, and the points are the averages (n = 6). **D-F.** Comparisons between impedance measurements using a Fluke PM6306 LCR meter and our TEEI-PSEP system. In F, the LCR meter plot is obscured by the overlapping TEEI-PSEP plot.

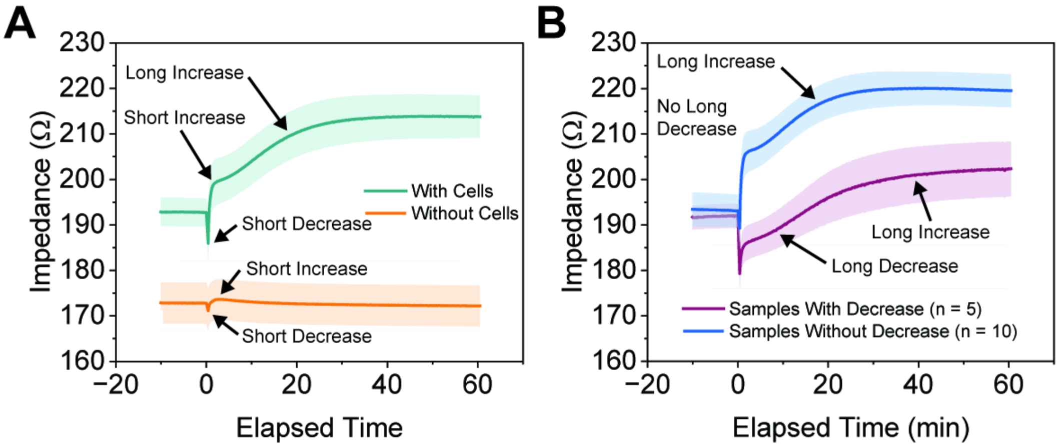


**Figure S2. Post-EP TEEI Increase.** Alternative representations of the data in Figure 2A. **A.** Impedance measurements with and without cells that were used to calculate the change in TEEI. The short increase and short decrease features are present with and without cells, suggesting they are not part of the post-EP TEEI increase. **B.** The impedance with cells separated into samples with the long decrease and samples without the long decrease. The long decrease is not present in most of the samples, which is why it is not present in the overall average with cells. The error bars in A and B represent the standard error of the mean (SEM) (n = 15).

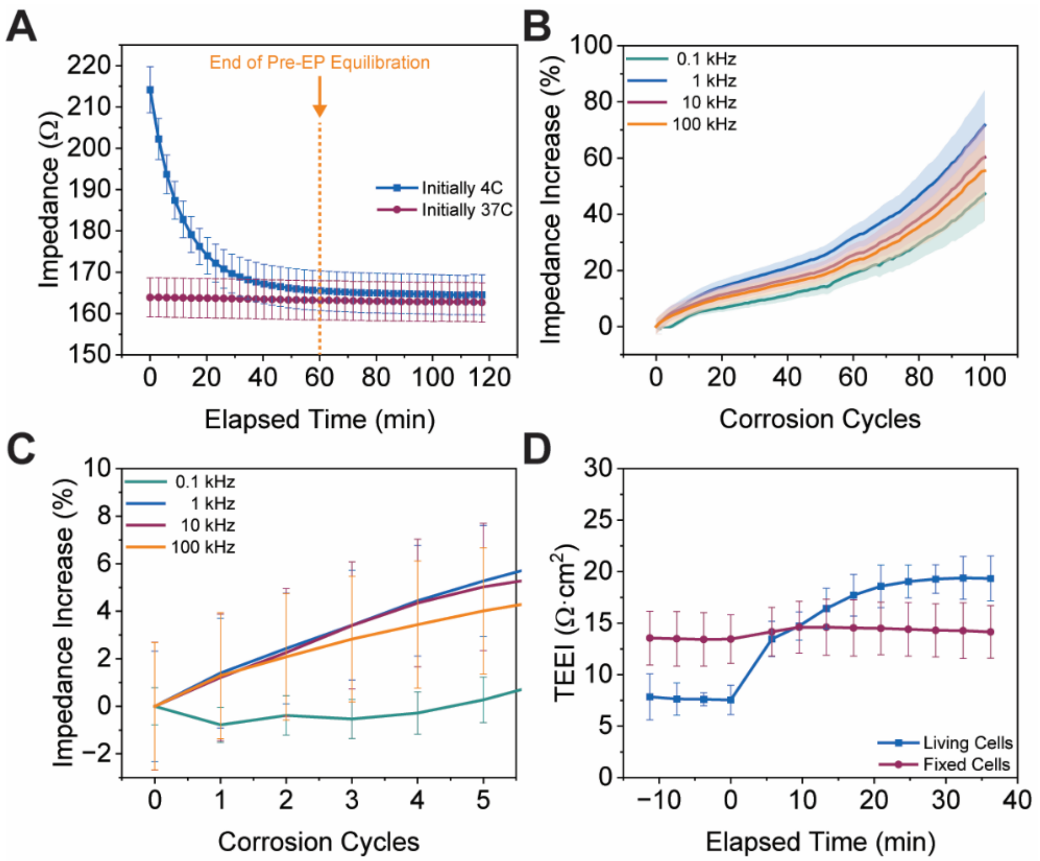


**Figure S3. Evaluation of Cell-Free and Passive Cell Hypotheses. A.** Impedance measurements for samples without cells starting at 37°C and starting at 4°C and placed in a 37°C incubator. The end of pre-EP equilibration denotes where electroporation would begin in a normal experiment. **B.** Impedance measurements after repeated corrosion cycles. **C.** Impedance measurements after repeated corrosion cycles showing the change between individual cycles. **D.** Post-EP TEEI measurements for living and fixed cells. The error bars represent the standard error of the mean (SEM) (n = 6).

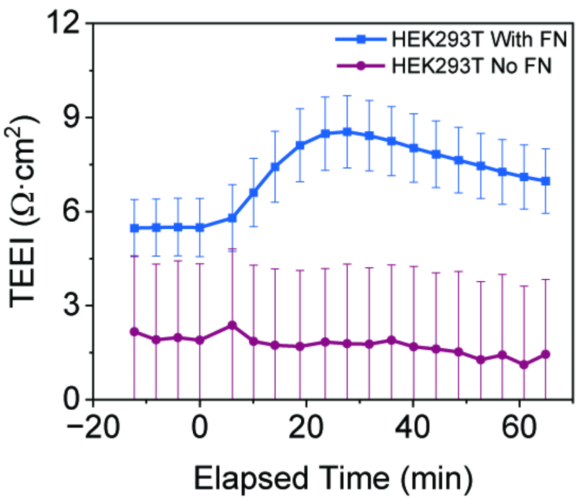


**Figure S4. The HEK293T Post-EP TEEI Response is Dependent on Fibronectin Coating.** TEEI measurements for HEK293T cells cultured with the standard FN coating and with no FN coating. The error bars represent the standard error of the mean (SEM) (n = 6).

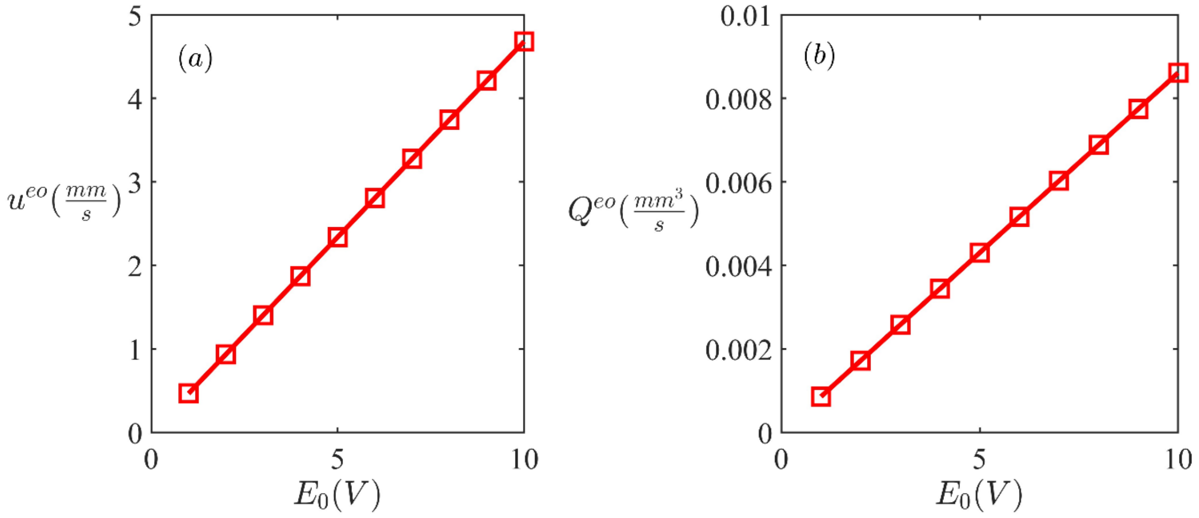


**Figure S5. Theoretical electro-osmosis velocity and flow rate resulting from electro-osmosis. A.** Electro-osmosis velocity as function of the voltage difference. The parameters used are $\zeta_w = -12\, mv$, $\varepsilon = 78.4\varepsilon_0 = 6.9415e-10$, $\eta = 8.9e-4$, and the channel length of $L = 20\mu m$. **B.** The time-averaged total flow rate across the substrate resulting from the electro-osmosis. Note that for this calculation, we account for the 2% cycle duty of the applied voltage and use a substrate with an area (A) of 0.336 $cm^2$ and a channel density of 2e6 (channel number/$cm^2$).

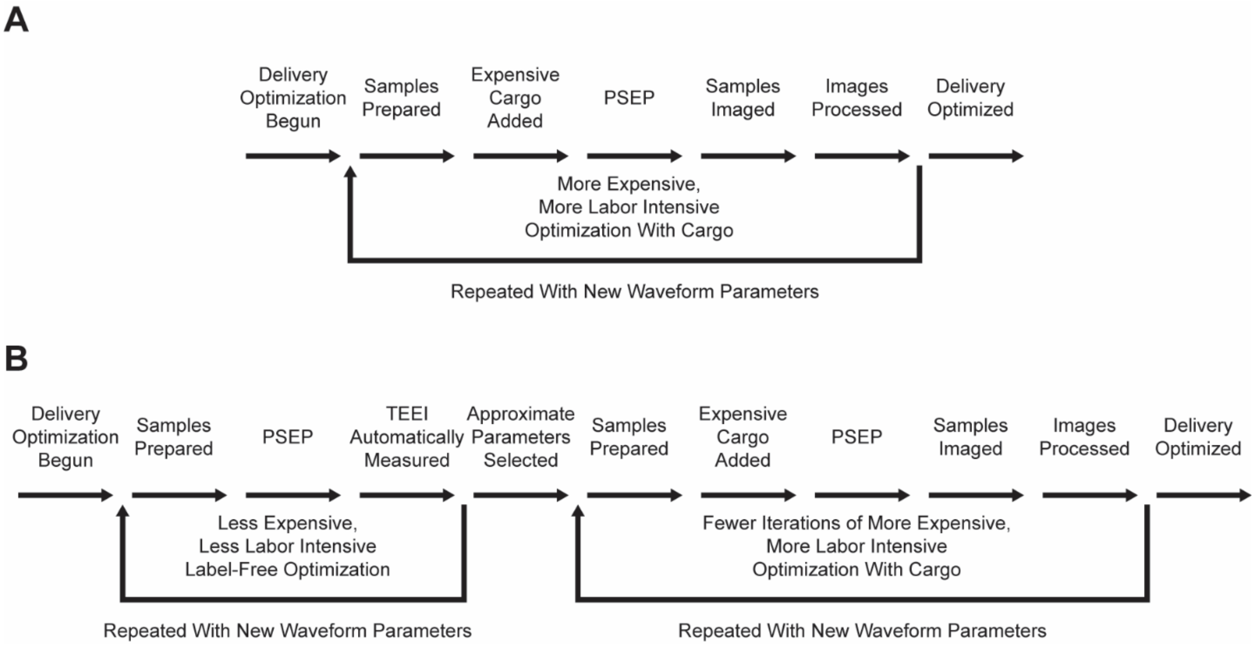


**Figure S6. PSEP Iterative Optimization.** Depictions of PSEP optimization with and without label-free delivery. **A.** Standard PSEP optimization process without label-free delivery requiring more expensive, more labor-intensive optimization cycles. **B.** Proposed PSEP optimization process with label-free delivery. Label-free delivery is used with an initial series of optimization cycles to obtain approximate parameters with reduced time and cost. PSEP is then optimized with the cargo of interest, but with fewer iterations because approximate parameters have already been found.

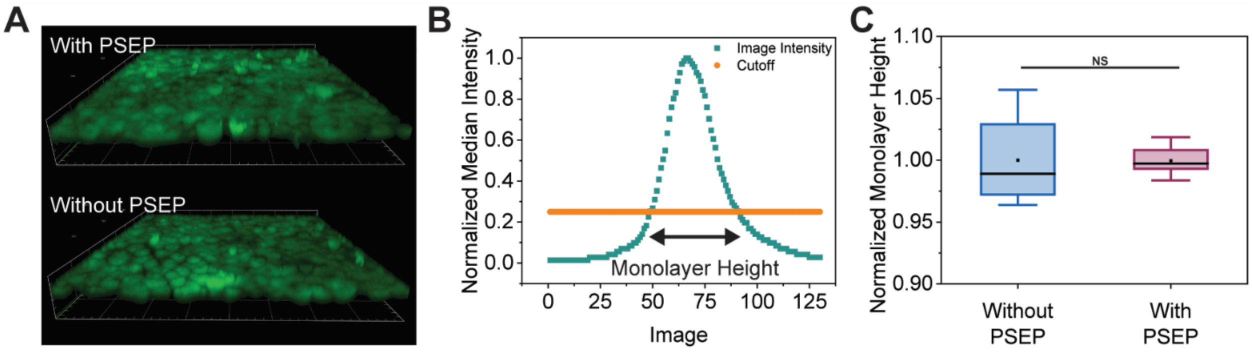


**Figure S7. No Change in Cell Monolayer Height Following PSEP. A.** 3D fluorescent Z-stacks of cell monolayers with and without PSEP labelled with calcein. **B.** Method for measuring cell monolayer height using the normalized median intensity of each image in the Z-stack and a cutoff of 0.25. **C.** Cell monolayer height measured using Z-stack with and without PSEP. The shaded areas represent the interquartile ranges (IQRs), the error bars represent 1.5 times the IQRs, the horizontal lines are the medians, and the points are the averages (n = 6).

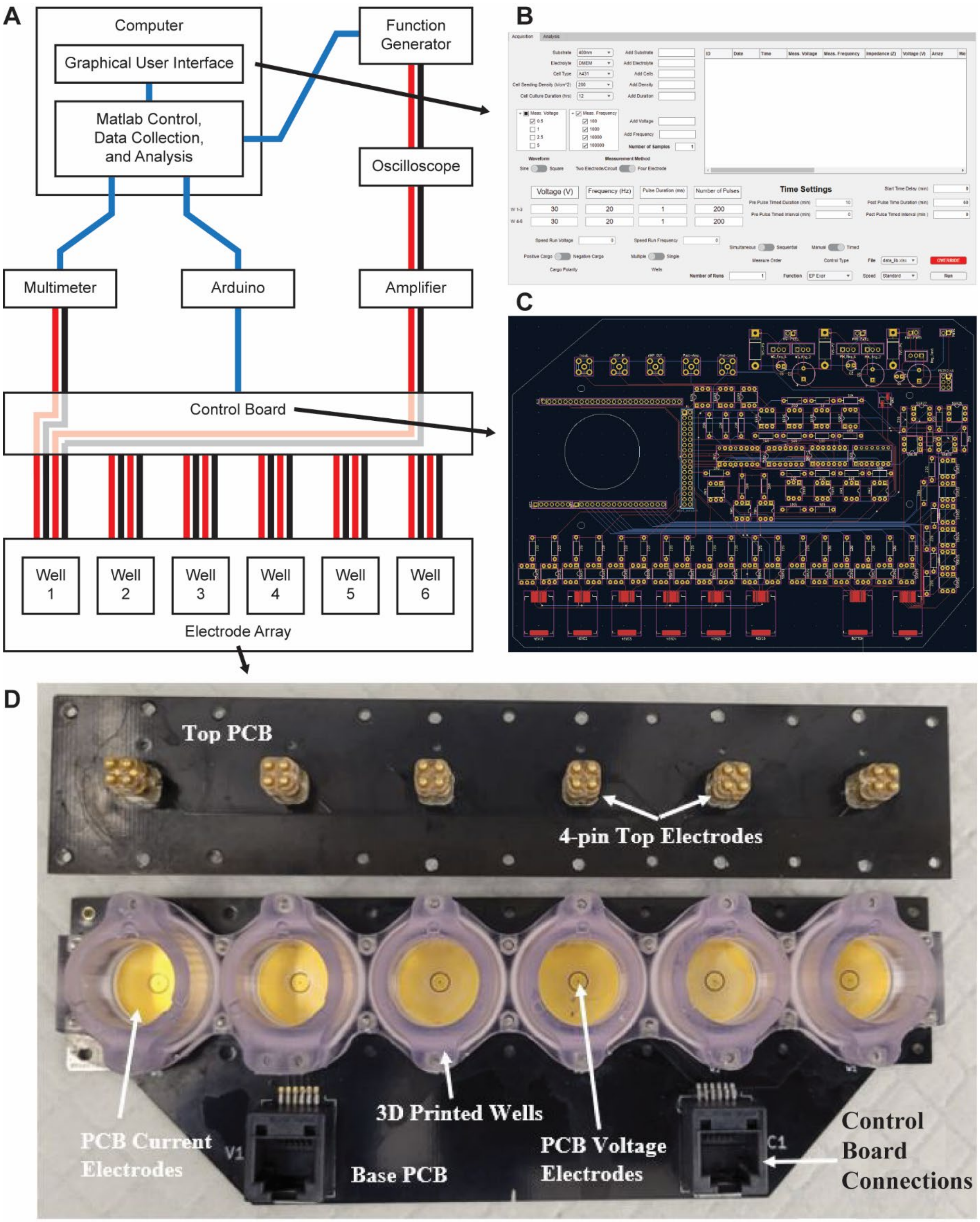


**Figure S8. Device Overview. A.** Simplified depiction of system components and connections. Blue lines represent data connections. Red lines and black lines represent positive and negative connections, respectively. In the current configuration, well 1 is being pulsed and measured. **B.** Graphical user interface. **C.** Control board. **D.** Electrode array.

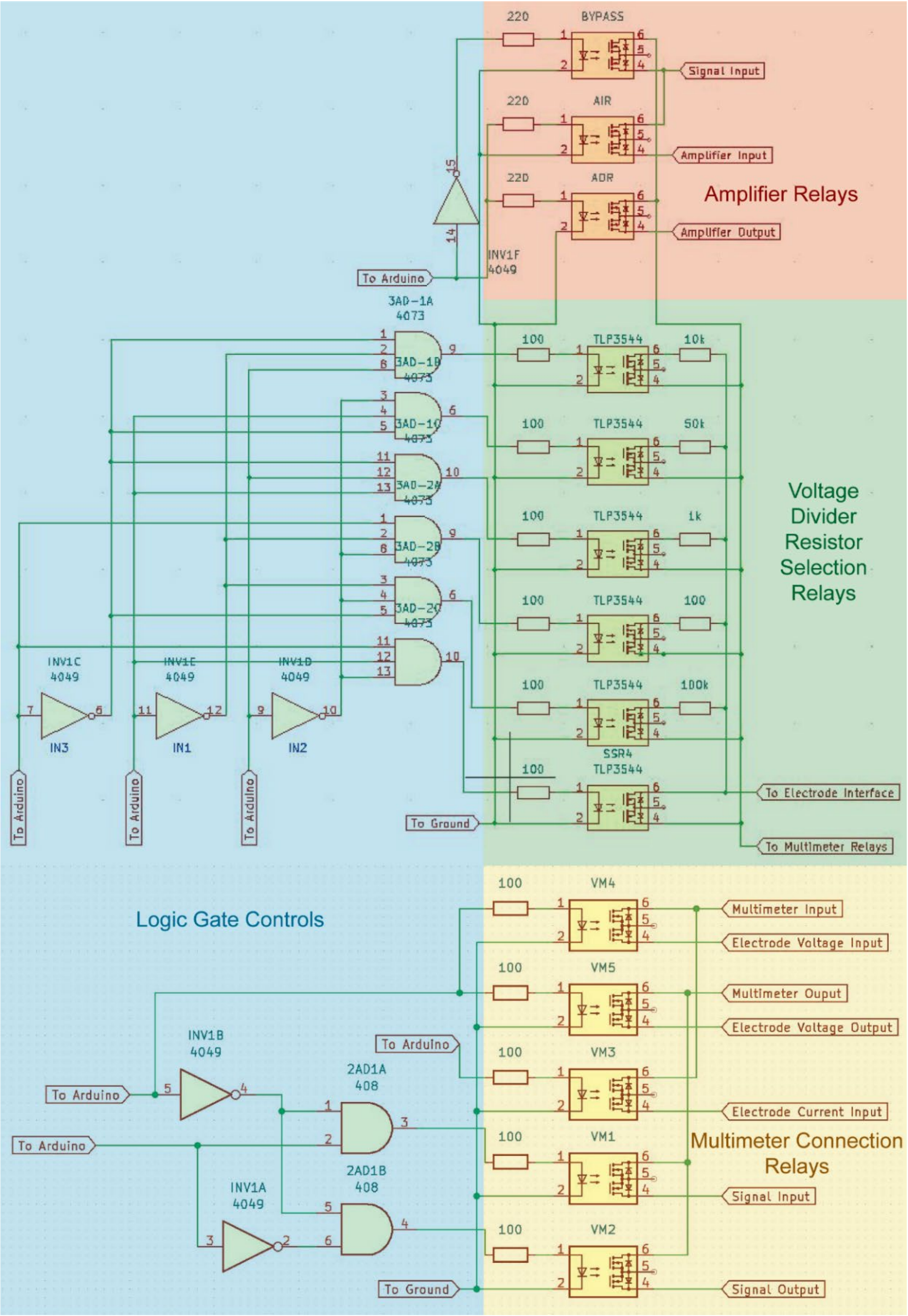


**Figure S9. Schematic of Notable Portions of the Control Board.** Notable portions of the control board including the logic gate controls in blue, amplifier relays in red, voltage divider resistor section relays in green, and multimeter connection relays in yellow.

### SUPPLEMENTAL TABLES

#### Table S1. Device Validation Using LCR Meter

| Configuration | Figure | Component | Component Value |
|---|---|---|---|
| Resistor | S1D | Resistor | 2 kΩ |
| Series Resistor and Capacitor | S1E | Resistor | 2 kΩ |
| | | Capacitor | 1 μF |
| Resistor in Series with Parallel Resistor and Capacitor | S1F | Series Resistor | 100 Ω |
| | | Parallel Resistor | 220 Ω |
| | | Capacitor | 1 μF |

### Table S2. Effective Measurement Rates

| Measurement Condition: 1 Well, 1 Frequency | | | Measurement Condition: 6 Wells, 1 Frequency | | | Measurement Condition: 6 Wells, 4 Frequencies | | |
|---|---|---|---|---|---|---|---|---|
| Well | Frequency | Measurement | Well | Frequency | Measurement | Well | Frequency | Measurement |
| 1 | 1 kHz | 1 | 1 | 1 kHz | 1 | 1 | 100 Hz | 1 |
| 1 | 1 kHz | 2 | 2 | 1 kHz | 1 | 2 | 100 Hz | 1 |
| 1 | 1 kHz | 3 | 3 | 1 kHz | 1 | 3 | 100 Hz | 1 |
| 1 | 1 kHz | 4 | 4 | 1 kHz | 1 | 4 | 100 Hz | 1 |
| 1 | 1 kHz | 5 | 5 | 1 kHz | 1 | 5 | 100 Hz | 1 |
| 1 | 1 kHz | 6 | 6 | 1 kHz | 1 | 6 | 100 Hz | 1 |
| 1 | 1 kHz | 7 | 1 | 1 kHz | 2 | 1 | 1 kHz | 1 |
| 1 | 1 kHz | 8 | 2 | 1 kHz | 2 | 2 | 1 kHz | 1 |
| 1 | 1 kHz | 9 | 3 | 1 kHz | 2 | 3 | 1 kHz | 1 |
| 1 | 1 kHz | 10 | 4 | 1 kHz | 2 | 4 | 1 kHz | 1 |
| 1 | 1 kHz | 11 | 5 | 1 kHz | 2 | 5 | 1 kHz | 1 |
| 1 | 1 kHz | 12 | 6 | 1 kHz | 2 | 6 | 1 kHz | 1 |
| 1 | 1 kHz | 13 | 1 | 1 kHz | 3 | 1 | 10 kHz | 1 |
| 1 | 1 kHz | 14 | 2 | 1 kHz | 3 | 2 | 10 kHz | 1 |
| 1 | 1 kHz | 15 | 3 | 1 kHz | 3 | 3 | 10 kHz | 1 |
| 1 | 1 kHz | 16 | 4 | 1 kHz | 3 | 4 | 10 kHz | 1 |
| 1 | 1 kHz | 17 | 5 | 1 kHz | 3 | 5 | 10 kHz | 1 |
| 1 | 1 kHz | 18 | 6 | 1 kHz | 3 | 6 | 10 kHz | 1 |
| 1 | 1 kHz | 19 | 1 | 1 kHz | 4 | 1 | 100 kHz | 1 |
| 1 | 1 kHz | 20 | 2 | 1 kHz | 4 | 2 | 100 kHz | 1 |
| 1 | 1 kHz | 21 | 3 | 1 kHz | 4 | 3 | 100 kHz | 1 |
| 1 | 1 kHz | 22 | 4 | 1 kHz | 4 | 4 | 100 kHz | 1 |
| 1 | 1 kHz | 23 | 5 | 1 kHz | 4 | 5 | 100 kHz | 1 |
| 1 | 1 kHz | 24 | 6 | 1 kHz | 4 | 6 | 100 kHz | 1 |
| Effective Measurement rate: 147 mHz | | | Effective Measurement rate: 25 mHz | | | Effective Measurement rate: 6 mHz | | |

Note: Most experiments used the 6 wells, 4 frequencies condition.

**Table S3. Cell Culture Parameters**

| Experiment | Figure | Seeding Density | Cell Line | Cell Culture Media | Fibronectin Concentration |
|---|---|---|---|---|---|
| Seeding Density | 2A | 50 k·cm$^{-2}$ | A431 | DMEM with 10% FBS and 1% P/S | 1 μg·mL$^{-1}$ |
| | | 100 k·cm$^{-2}$ | | | |
| | | 150 k·cm$^{-2}$ | | | |
| | | 200 k·cm$^{-2}$ | | | |
| Cell Lines | 2B | 200 k·cm$^{-2}$ | A431 | DMEM with 10% FBS and 1% P/S | 1 μg·mL$^{-1}$ |
| | | 400 k·cm$^{-2}$ | HEK293T | | |
| Fibronectin Concentration | 2C | 200 k·cm$^{-2}$ | A431 | DMEM with 10% FBS and 1% P/S | 0 μg·mL$^{-1}$ |
| | | | | | 0.1 μg·mL$^{-1}$ |
| | | | | | 1 μg·mL$^{-1}$ |
| | | | | | 10 μg·mL$^{-1}$ |

Areas marked in gray indicate standard cell culture parameters. All cell culture experiments used the standard substrates and electroporation waveforms as described in the materials and methods section.

**Table S4. Electroporation Waveform Parameters**

| Experiment | Figure | Voltage | Pulse Duration | Pulse Frequency | Pulse Number | Train Number | Train Frequency |
|---|---|---|---|---|---|---|---|
| Voltage | 3B | 20 V<br>30 V<br>34 V<br>38 V | 1 ms | 20 Hz | 200 pulses/ train | 1 train | N/A |
| Pulse Duration | 3C | 30 V | 0.3 ms<br>1 ms<br>3 ms | 20 Hz | 200 pulses/ train | 1 train | N/A |
| Pulse Frequency | 3D | 30 V | 1 ms | 5 Hz<br>20 Hz<br>80 Hz | 200 pulses/ train | 1 train | N/A |
| Pulse Number | 3E | 30 V | 1 ms | 20 Hz | 50 pulses/ train<br>200 pulses/ train<br>800 pulses/ train | 1 train | N/A |

Areas marked in gray indicate standard waveform parameters. All waveform experiments used the standard cell culture conditions as described in the materials and methods section.

### Table S5. Electroporation Impedance Measurement Studies

| Electroporation System | | | | | Electroporation Waveform | | | | | Response | | | | | Ref. |
|---|---|---|---|---|---|---|---|---|---|---|---|---|---|---|---|
| **Vivo/ Vitro** | **Cell Scale** | **Cell Line** | **Electrode Format** | **Number of Electrodes per Sample** | **Voltage or Electric Field** | **Pulse Duration** | **Pulse Frequency** | **Number of Pulses** | **Shape** | **Impedance Change** | **% Change** | **Recovery Time** | **Frequencies Measured** | **Sampling Rate** | |
| Vitro | Tissue | human stratum corneum | | 4 | 50- 250 V | | 0.1 Hz- 93 µHz | 1-3 | Exponential | Decrease | > 70% | < 1 min | 1 kHz- 100 kHz | 94- 1500 Hz | (*7*) |
| Vivo | Tissue | rat muscle cells, rat liver cells | Plates | 2 | 50- 550 V | 100 µs | 1 Hz | 8 | Square | Decrease | < 40% | | | 25- 100 MHz | (*8*) |
| Vivo | Tissue | rat liver cells | Plates | 4 | 450- 1500 V/cm | 100 µs | 10 Hz | 8 | | Increase | < 15% | ~ 7 min | 1 kHz- 400 kHz | | (*9*) |
| Vivo | Tissue | rat liver cells | Rods | 18 (2 EP, 16 measurement) | 450- 1500 V/cm | 100 µs | 10 Hz | 8 | | Decrease | < 45% | | | | (*10*) |
| Vivo | Tissue | C57 B1/6 mouse sarcoma cells | Plates | 4 | 450- 3500 V/cm | 100- 1000 µs | 0.03- 10 Hz | 8-80 | Square | Decrease | < 80% | > 30 min | 1 kHz- 400 kHz | 500 Hz | (*11*) |
| Vitro | Double Monolayer | rat brain microvascular endothelial cells, cerebral astrocytes | | | 100- 400 kV/m | | 1 Hz | 100-400 | Exponential | Decrease | < 10% | > 15 hrs | | 0.28 mHz | (*12*) |
| Vitro | Monolayer | NRK, HEK 293, CHO, NIH-3T3, Hep G2 | PCB | 2 | 3-5 $V_{rms}$ | 25 µs | 40 kHz | 8000- 20000 | Sinusoidal | Decrease then increase | < 30% dec., < 20% inc. | ~ 30 min | 4 kHz | 1 Hz | (*13*) |
| Vitro | Monolayer | CHO, 3T3-L1 | PCB | 4 | 200- 1400 V/cm | | | 8 | Square biphasic | Decrease | < 20% | | | | (*14*) |
| Vitro | Monolayer | C2C12 | PCB | 4 | 400- 1200 V/cm | 100 µs | 1 Hz | 8 | Square biphasic | Decrease | < 35 % | > 1 sec | 5-44 kHz | 1 kHz | (*15*) |
| Vitro | Monolayer | HeLa | PCB | | 10-14 V | 100 µs | 1 Hz | 10 | Square | Decrease | < 60% | < 10 min | 100 kHz | | (*16*) |
| Vitro | Monolayer | DC3F, C2C12, HeLa | PCB | 4 | 600- 1400 V/cm | 100 µs | 1 Hz | 8 | Square biphasic | Decrease | < 25% | > 8 sec | 7 kHz- 1.12 MHz | 1 kHz | (*17*) |
| Vitro | Monolayer | NRK, HEK 293, Hep G2, CHO | PCB | | 2-7 (5.0) V | 200, 500 ms | 40 kHz | | Sine | Decrease | 30-50% | 1-3.5+ hrs | 4 kHz | | (*18*) |
| Vitro | Monolayer | MCF-7 | | | 2.5-5 V | 1 µs | | 0-200 | Square | Decrease | < 50% | 2 hrs | | 3 mHz | (*19*) |
| Vitro | Suspended Pellet | Rabbit erythrocytes | | | 240-1140 V, 2300 V/cm | 10 us- 40 µs | N/A | 1 | | Decrease, increase | > 90% dec. < 350% inc. | ~ 40 min | | | (*20*) |
| Vitro | Adherent Single Cell | GH3, Jurkat, PC-12, HeLa | Wires | 2 | 12000 V/cm | 60 ns | N/A | 1 | Square | Decrease | < 75% | > 15 min | | | (*21*) |
| Vitro | Adherent Single Cell | Primary cardiomyocytes | PSEP | | 3 V | 200 µs | 20 Hz | 20 | Square | Decrease | | < 102 min | | 15 kHz | (*22*) |
| Vitro | Suspended Single Cell | HeLa, CHO-K1 | PCB | 2 | 8 V | | 50 kHz | | | Decrease | | | 20 kHz- 20 MHz | | (*23*) |

This table is intended to provide an overview of electroporation impedance studies that have been performed but is not exhaustive.

**Table S6. Corrosion**

| Frequency | 100 Hz | | 1 kHz | | 10 kHz | | 100 kHz | |
|---|---|---|---|---|---|---|---|---|
| Pulses | Average Z (ΔZ%) | SEM (n=6) | Average Z (ΔZ%) | SEM (n=6) | Average Z (ΔZ%) | SEM (n=6) | Average Z (ΔZ%) | SEM (n=6) |
| 0 | 288.81 | 2.2513 | 169.52 | 3.9447 | 155.50 | 4.1862 | 153.51 | 4.1217 |
| 5 | 289.60 (+0%) | 2.7713 | 177.302 (+5%) | 3.9946 | 163.03 (+5%) | 4.1416 | 159.74 (+4%) | 4.0673 |
| 10 | 299.86 (+4%) | 2.7469 | 183.66 (+8%) | 4.0069 | 167.01 (+7%) | 4.1434 | 163.75 (+7%) | 4.0009 |
| **20** | **307.76 (+7%)** | **4.0366** | **192.43 (+14%)** | **4.3298** | **173.4 (+12%)** | **4.3565** | **169.41 (+10%)** | **4.2452** |
| 50 | 329.57 (+14%) | 7.4060 | 211.28 (25%) | 5.4929 | 186.58 (+20%) | 4.9851 | 181.65 (+18%) | 4.7313 |
| 100 | 425.32 (+47%) | 27.660 | 290.7 (+71%) | 20.825 | 251.51 (+62%) | 18.464 | 241.29 (57%) | 17.218 |

Note: Electrodes were replaced after 20 pulses to minimize variation in impedance measurements.

**Movie S1.**

A timelapse of motion within the A431 cell monolayer before electroporation. Images were taken every minute for 10 minutes. The cells were genetically engineered to express green fluorescent protein (GFP) conjugated epithelial cadherin (E-cadherin).

**Movie S2.**

A timelapse of motion within the same A431 cell monolayer as Movie S1 immediately after electroporation. Images were taken every minute for 10 minutes.

### References


1. J. R. Brooks, I. Mungloo, S. Mirfendereski, J. P. Quint, D. Paul, A. Jaberi, J. S. Park, R. Yang, An equivalent circuit model for localized electroporation on porous substrates. *Biosensors and Bioelectronics* **199**, 113862 (2022).
2. J. Seebach, P. Dieterich, F. Luo, H. Schillers, D. Vestweber, H. Oberleithner, H.-J. Galla, H.-J. Schnittler, Endothelial Barrier Function under Laminar Fluid Shear Stress. *Laboratory Investigation* **80**, 1819-1831 (2000).
3. J. P. Vigh, A. Kincses, B. Ozgür, F. R. Walter, A. R. Santa-Maria, S. Valkai, M. Vastag, W. Neuhaus, B. Brodin, A. Dér, M. A. Deli, Transendothelial Electrical Resistance Measurement across the Blood–Brain Barrier: A Critical Review of Methods. *Micromachines* **12**, 10.3390/mi12060685 (2021).
4. B. Srinivasan, A. R. Kolli, M. B. Esch, H. E. Abaci, M. L. Shuler, J. J. Hickman, TEER Measurement Techniques for In Vitro Barrier Model Systems. *Journal of Laboratory Automation* **20**, 107-126 (2015).
5. S.-E. Choi, H. Khoo, S. C. Hur, Recent Advances in Microscale Electroporation. *Chemical Reviews* **122**, 11247-11286 (2022).
6. J. L. Anderson, Colloid Transport by Interfacial Forces. *Annual Review of Fluid Mechanics* **21**, 61-99 (1989).
7. U. Pliquett, R. Langer, J. C. Weaver, Changes in the passive electrical properties of human stratum corneum due to electroporation. *Biochimica et Biophysica Acta (BBA) - Biomembranes* **1239**, 111-121 (1995).
8. D. Cukjati, D. Batiuskaite, F. André, D. Miklavčič, L. M. Mir, Real time electroporation control for accurate and safe in vivo non-viral gene therapy. *Bioelectrochemistry* **70**, 501-507 (2007).
9. A. Ivorra, B. Rubinsky, In vivo electrical impedance measurements during and after electroporation of rat liver. *Bioelectrochemistry* **70**, 287-295 (2007).
10. Y. Granot, B. Rubinsky, Methods of optimization of electrical impedance tomography for imaging tissue electroporation. *Physiological Measurement* **28**, 1135 (2007).
11. A. Ivorra, B. Al-Sakere, B. Rubinsky, L. M. Mir, In vivo electrical conductivity measurements during and after tumor electroporation: conductivity changes reflect the treatment outcome. *Physics in Medicine & Biology* **54**, 5949 (2009).
12. Z. J. Xing, D. G. Rong, Z. Jie, Z. Y. Chun, Z. Y. Jun, G. G. Zhen, Detrimental Effect of Electromagnetic Pulse Exposure on Permeability of In Vitro Blood-brain-barrier Model. *Biomedical and Environmental Sciences* **26**, 128 (2013).
13. J. A. Stolwijk, C. Hartmann, P. Balani, S. Albermann, C. R. Keese, I. Giaever, J. Wegener, Impedance analysis of adherent cells after in situ electroporation: Non-invasive monitoring during intracellular manipulations. *Biosensors and Bioelectronics* **26**, 4720-4727 (2011).
14. T. García-Sánchez, M. Guitart, J. Rosell-Ferrer, A. M. Gómez-Foix, R. Bragós, A new spiral microelectrode assembly for electroporation and impedance measurements of adherent cell monolayers. *Biomedical Microdevices* **16**, 575-590 (2014).
15. T. García-Sánchez, A. Azan, I. Leray, J. Rosell-Ferrer, R. Bragós, L. M. Mir, Interpulse multifrequency electrical impedance measurements during electroporation of adherent differentiated myotubes. *Bioelectrochemistry* **105**, 123-135 (2015).
16. X. Guo, R. Zhu, Controllable in-situ cell electroporation with cell positioning and impedance monitoring using micro electrode array. *Scientific Reports* **6**, 31392 (2016).

17. T. García-Sánchez, R. Bragós, L. M. Mir, In vitro analysis of various cell lines responses to electroporative electric pulses by means of electrical impedance spectroscopy. *Biosensors and Bioelectronics* **117**, 207-216 (2018).
18. J. A. Stolwijk, J. Wegener, Impedance analysis of adherent cells after in situ electroporation-mediated delivery of bioactive proteins, DNA and nanoparticles in µL-volumes. *Scientific Reports* **10**, 21331 (2020).
19. D. Lee, S. S. Y. Chan, N. Aksic, N. Bajalovic, D. K. Loke, Ultralong-Time Recovery and Low-Voltage Electroporation for Biological Cell Monitoring Enabled by a Microsized Multipulse Framework. *ACS Omega* **6**, 35325-35333 (2021).
20. I. G. Abidor, L. H. Li, S. W. Hui, Studies of cell pellets: II. Osmotic properties, electroporation, and related phenomena: membrane interactions. *Biophysical Journal* **67**, 427-435 (1994).
21. A. G. Pakhomov, J. F. Kolb, J. A. White, R. P. Joshi, S. Xiao, K. H. Schoenbach, Long-lasting plasma membrane permeabilization in mammalian cells by nanosecond pulsed electric field (nsPEF). *Bioelectromagnetics* **28**, 655-663 (2007).
22. D. Xu, J. Fang, H. Wang, X. Wei, J. Yang, H. Li, T. Yang, Y. Li, C. Liu, N. Hu, Scalable Nanotrap Matrix Enhanced Electroporation for Intracellular Recording of Action Potential. *Nano Letters* **22**, 7467-7476 (2022).
23. S. C. Bürgel, C. Escobedo, N. Haandbæk, A. Hierlemann, On-chip electroporation and impedance spectroscopy of single-cells. *Sensors and Actuators B: Chemical* **210**, 82-90 (2015).
